# Identification of uterine pacemaker regions at the myometrial-placental interface

**DOI:** 10.1101/152678

**Authors:** E. Josiah Lutton, Wim J. E. P. Lammers, Sean James, Hugo A. van den Berg, Andrew M. Blanks

## Abstract

Coordinated uterine contractions at the end of gestation are essential for delivering viable offspring in mammals. Contractions are initiated by an electrical signal at the plasma membrane of uterine muscle cells, leading to voltage-dependent calcium entry, and subsequent activation of the intracellular contractile machinery. In contrast to other visceral smooth muscles, it is not known where excitation within the uterus is initiated, and no defined pacemaking region has hitherto been identified. Using a combination of multi-electrode array recordings and high-resolution computational reconstruction of the three-dimensional micro-structure of late pregnant rat uterus, we demonstrate that electrical potentials are initiated in distinct structures within the placental bed of individual implantation sites. These previously unidentified structures represent modified smooth muscle bundles that are derived from bridges between the longitudinal and circular layers. Coordinated implantation and encapsulation by invading trophoblast give rise to isolated placental/myometrial interface bundles that directly connect to the overlying longitudinal smooth muscle layer. Furthermore, the numerous bridge structures co-localise with the vascular network located between the longitudinal and circular layers. Taken together, these observations imply that the anatomical structure of the uterus, combined with site-specific implantation, gives rise to emergent patterns of electrical activity that drive effective contractility during parturition. The identification of the pacemaking zones of the uterus has important consequences for the treatment of disorders of parturition such as preterm labor, postpartum hemorrhage and uterine dystocia.

## Introduction

Myometrial smooth muscle is capable of generating phasic contractions in the absence of stimuli from the central nervous system or circulating hormones [16]. As in all visceral muscles, contractions require the generation and propagation of electrical signals at the plasma membrane of cells, which are arranged in an electrotonically connected syncytium [15]. Tissues such as the heart and the stomach orchestrate electrical activity via dedicated anatomical structures that generate pacemaking potentials [29]. In contrast, the small intestine and bladder generate excitatory potentials at distinct but anatomically variable sites throughout the tissue [22, 27]. To date, no pacemaker regions with specific anatomical features have been described in the mammalian uterus, which is perhaps surprising given that coordinated forceful contractions are essential to parturition. Multi-electrode recordings of rat uteri demonstrated that potentials tend to initiate either at implantation sites on the mesometrial border, or near the ovarian end [28].

To shed light on the processes underlying activation in the rat myometrium, we established a three-step procedure that combines isochronal analysis of multi-electrode recordings with scans of histological slides of the same tissue specimens and automated image processing based on detection of cell nuclei, followed by 3D tissue reconstruction. This analysis enabled the correlation of the kinematics of reconstructed wavefronts of electrical activity propagating along the myometrium with a detailed reconstruction of tissue microarchitecture, at an unprecedented resolution level of 10 *μ*m for an entire organ measuring ~5 cm in length [30]. Using this strategy, we identified a defined histological structure in which electrical potentials are triggered by the integration of fetal and maternal stimuli, the “myometrial-placental pacemaker zone” (MPPZ).

## Results

In order to correlate electrical activity with anatomical structure, we first recorded electrical potentials from the serosal surface of rat myometrium as described previously [28]. After recording three representative one-minute time series of spontaneous activity per preparation, each taken from a different animal, the tissues were processed to capture the 3-dimensional histological microarchitecture as described previously [30]. Histological structure and electrical activity were subsequently collated to identify the anatomical sites where electrical excitation originates (Fig. 1).

**Figure 1.**
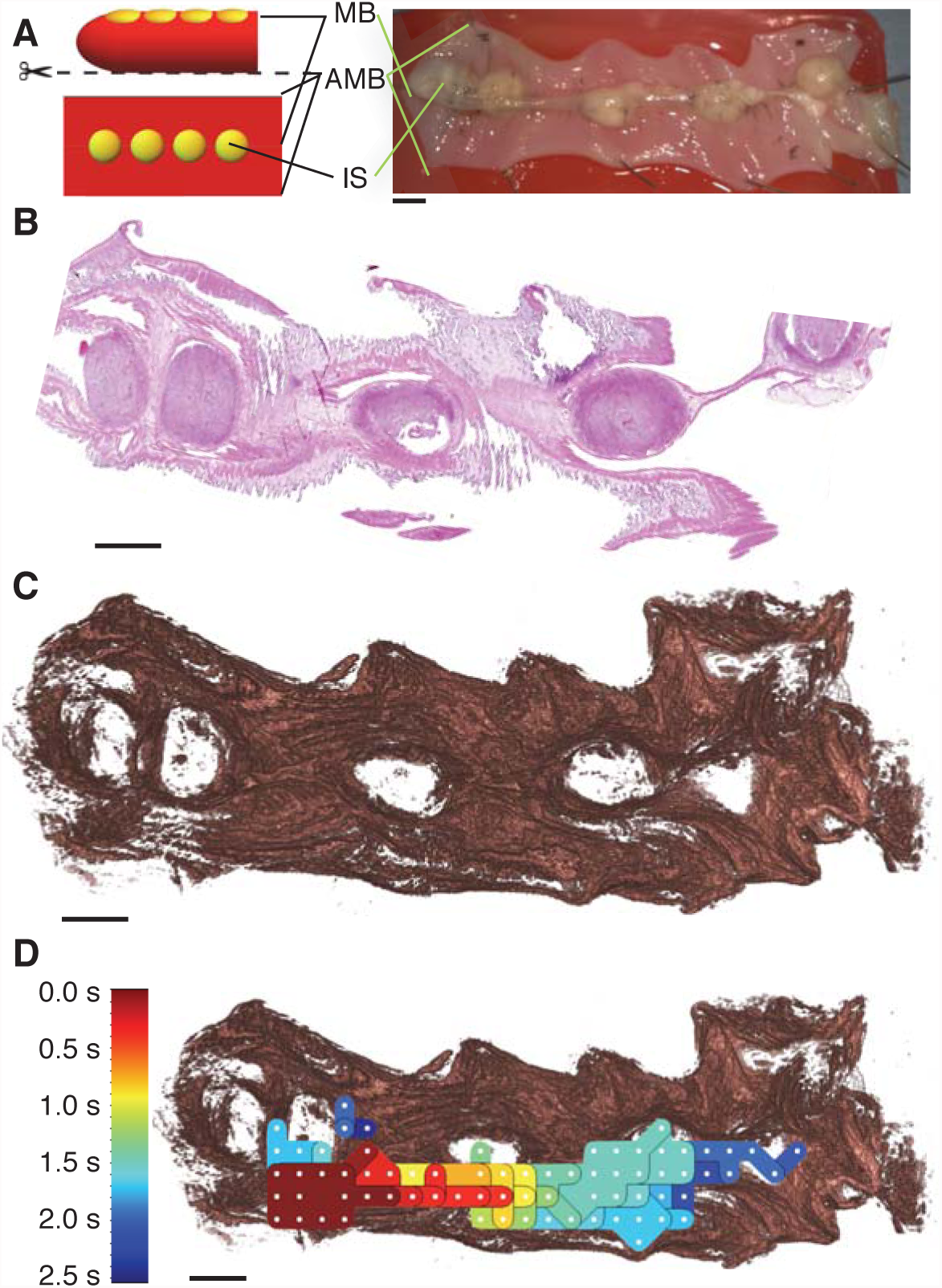
**A:** Three rat uteri were cut along the anti-mesometrial border (AMB) and pinned with the serosal side facing upwards, positioning the mesometrial border (MB), along with the implantation sites (IS), in the centre of the tissue. **B:** Each uterus was fixed in formalin, embedded in paraffin, sectioned into 5 *μ*m serial sections, and stained with haematoxylin and eosin. **C:** Detailed *in silico* reconstructions of the uteri were generated from these serial sections using a semi-automated image analysis pipeline. **D:** Multi-electrode array recordings were performed prior to fixation, which were processed to generate isochrone maps that represent the spread of electrical activity. These isochrone maps were compared to the reconstructed tissue to identify structural features in the tissue that affect the initiation and termination of the excitation waves. Colours in the isochrone map correspond to the time at which the excitation wave reaches the given area in the tissue (colour key on the left). All images representing tissue have the ovarian end of the tissue on the left. Scale bars represent 5 mm.

### Uterine Structure

The general higher-order structure of all samples analysed was consistent with that of a highly ordered inner circular layer of myometrial smooth muscle, which surrounds the decidualized stromal cells lining the lumen. The outer, sub-serosal, layer of the myometrial smooth muscle has a longitudinal orientation and is separated from the inner circular layer by connective tissue and vasculature.

In rodents, this general structure is well described, and is patterned during early post-natal development [4]. *In silico* tissue reconstruction visualises and charts 3-dimensional fibrous structure at high resolution [30], allowing us to discover novel anatomical features of the myometrium of functional significance. Detailed analysis revealed bridge-like structures that formed direct connections between the longitudinal and circular layers of the myometrium throughout the tissue (Fig. 2). Bridges were invariably associated with the vascular bed interspersed between the longitudinal and circular layer (Fig. 2**C–E**).

**Figure 2.**
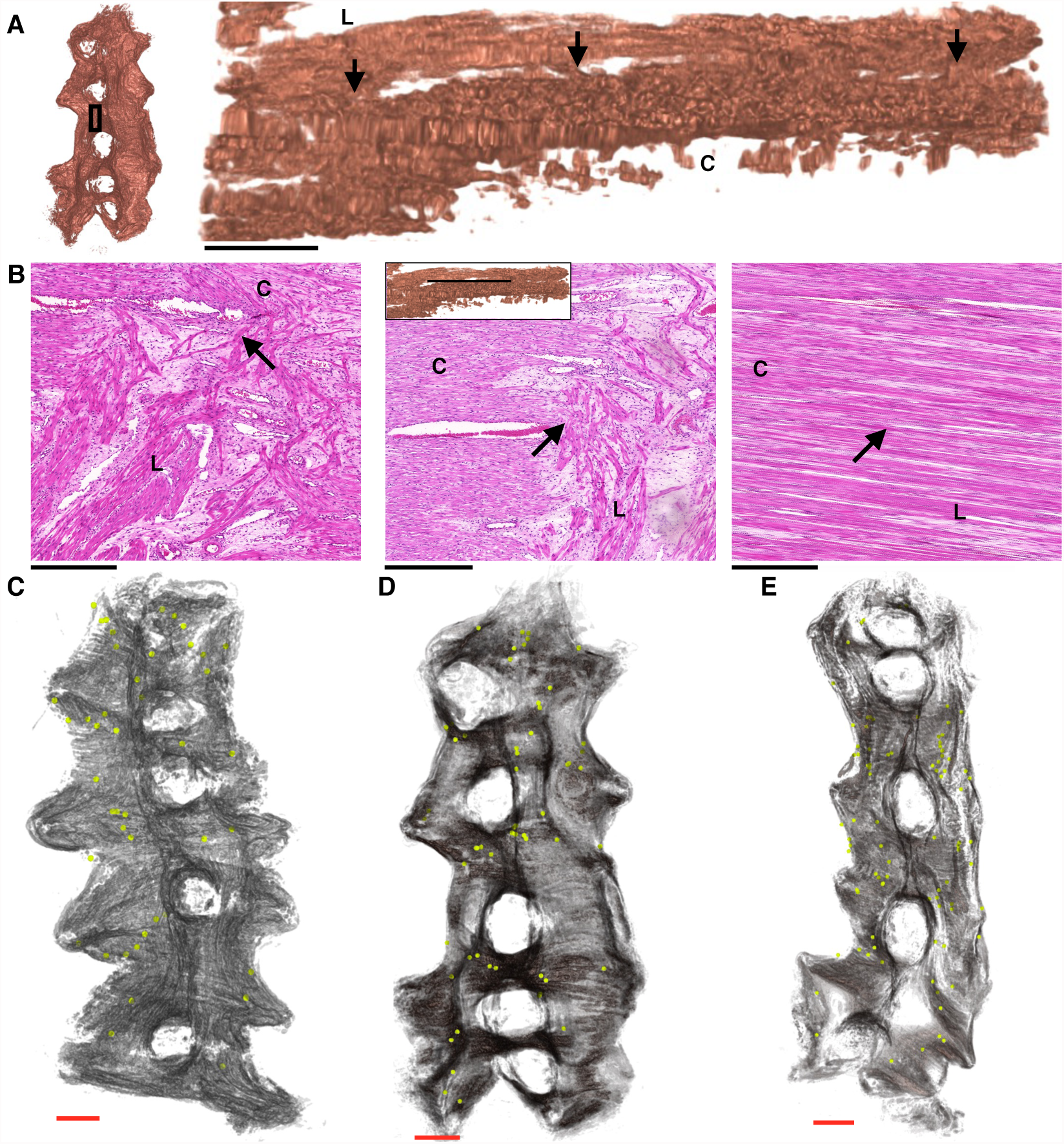
**A:** The three-dimensional reconstruction of the tissue in close proximity to the mesometrial border observed from a lateral viewpoint shows bridges joining the circular and longitudinal myometrium (arrows). **B:** These bridges can also be observed in the histological sections (haemotoxylin and eosin), appearing at multiple points along the length of **A. C–E:** locations of bridges (yellow) observed in a representative sample of histological slides. This demonstrates that these bridges are not exclusive to the mesometrial border. Subsequent analysis of fully resolved bridges revealed all structures were adjacent to vasculature. L: longitudinal myometrium; C: circular myometrium. Black scale bars represent 500 *μ*m, red scale bars represent 5 mm.

### Electrode array recordings and identification of myometrial placental pacemaker zones

We analyzed the spatio-temporal electrode array data in combination with the anatomical data to determine where electrical activity was originating. The electrode-array data were processed using a bespoke algorithm (see materials and methods), which identified similarities in the waveform of a train of action potentials that were used to cross-correlate signals at different locations in the array, allowing a given activation event to be tracked in time and space. Figure 3 presents the results as isochrone maps, superimposed on *in silico* reconstructions of the uteri. For each activation event, the electrode in closest proximity to the putative initiation point was identified and the local anatomical area was examined for common features that were confirmed for direct connectivity. In eight of the nine recordings, excitation was initiated in close proximity to an implantation site, with subsequent activity spreading along the mesometrial border. The final recording in the third tissue sample (bottom right in Figure 3) exhibited a different pattern of excitation, with activity initiated at the ovarian end of the tissue and spreading away from the mesometrial border; such excitations, though rarer than placental bed events, have been observed previously in these preparations [28]. Investigation of the histological slides at these initiation points revealed myometrial bundles in the placenta that were contiguous with the longitudinal layer (Fig 4). High-resolution 3-dimensional reconstructions of these structures revealed finger-like projections of myometrium in close proximity to placenta. The myometrial smooth muscle cells within these regions were distinguished by their stellate morphology from cells within the same bundle but situated outside of the placenta. These bundles appear to be modified bridge-like structures that have been captured by the invading placenta. Combination of the electrical recordings with the anatomical structure of the MPPZs and surrounding myometrial smooth muscle cell network allowed reconstruction of each event, demonstrating a distinct path of excitation from the MPPZ to the longitudinal fibres (Fig 5).

**Figure 3.**
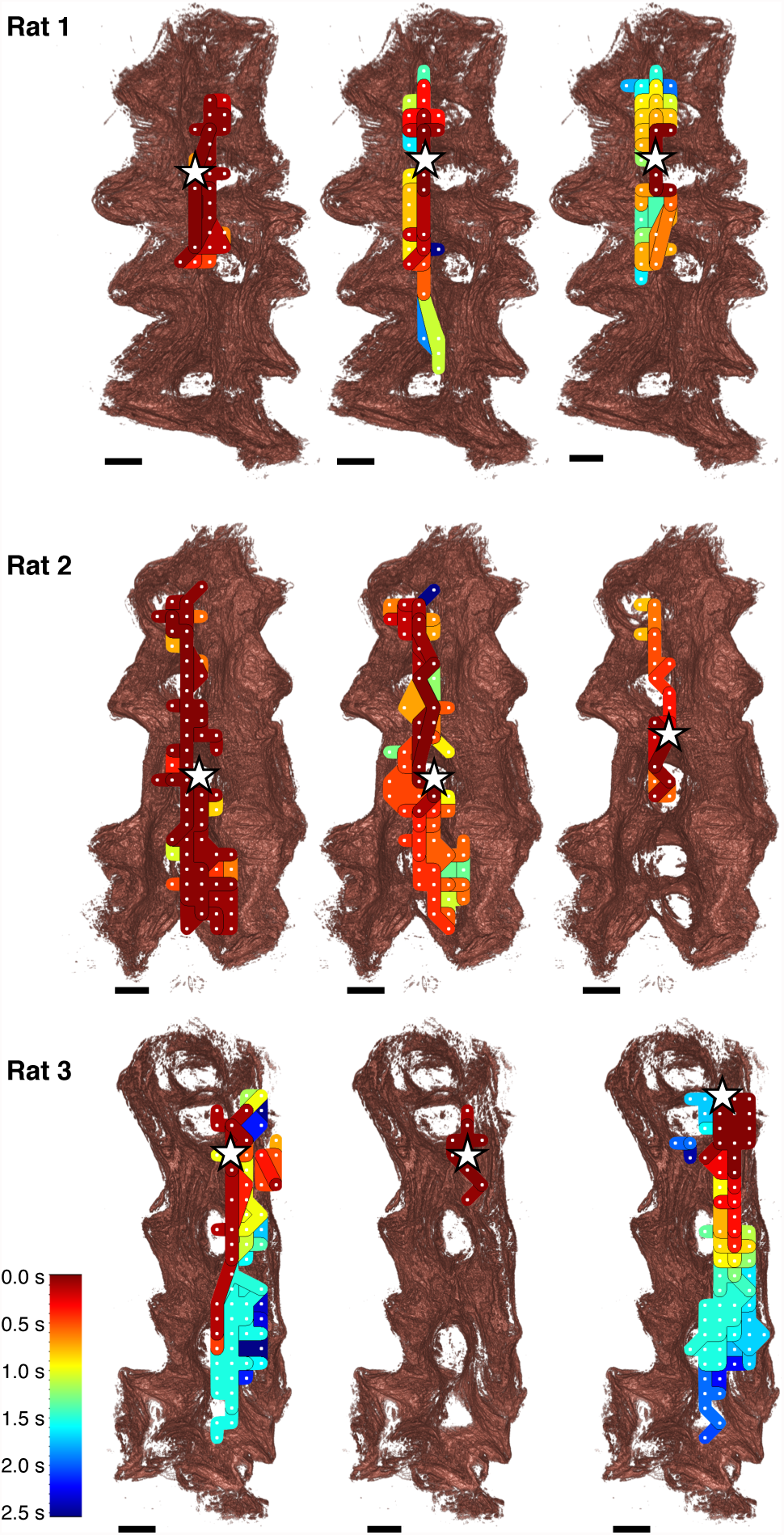
Isochrone maps generated from the multi-electrode array recordings in each of the uteri, superimposed onto the *in silico* reconstructions of the tissue. Each row shows isochrone maps of recordings taken from each of the rat uteri, where each recording covers one minute of activity, and consecutive recordings are separated by ~10 minutes. Isochrones are coloured based on the time that the excitation wave reached the given area according to the key shown, with each isochrone representing a 200ms time band. The electrode where the excitation wave was initiated in each recording was identified by eye and is indicated by a white star. Initiation points all lie along the mesometrial border in close proximity to the implantation sites. Excitation waves propagate along the mesometrial border in all but one recording (exception shown bottom-right). All recordings show excitation confined to one side of or along the mesometrial border, with many exhibiting excitation spreading across the implantation sites. Uteri are shown here with the ovarian end at the top of the image. Scale bars represent 5 mm.

**Figure 4.**
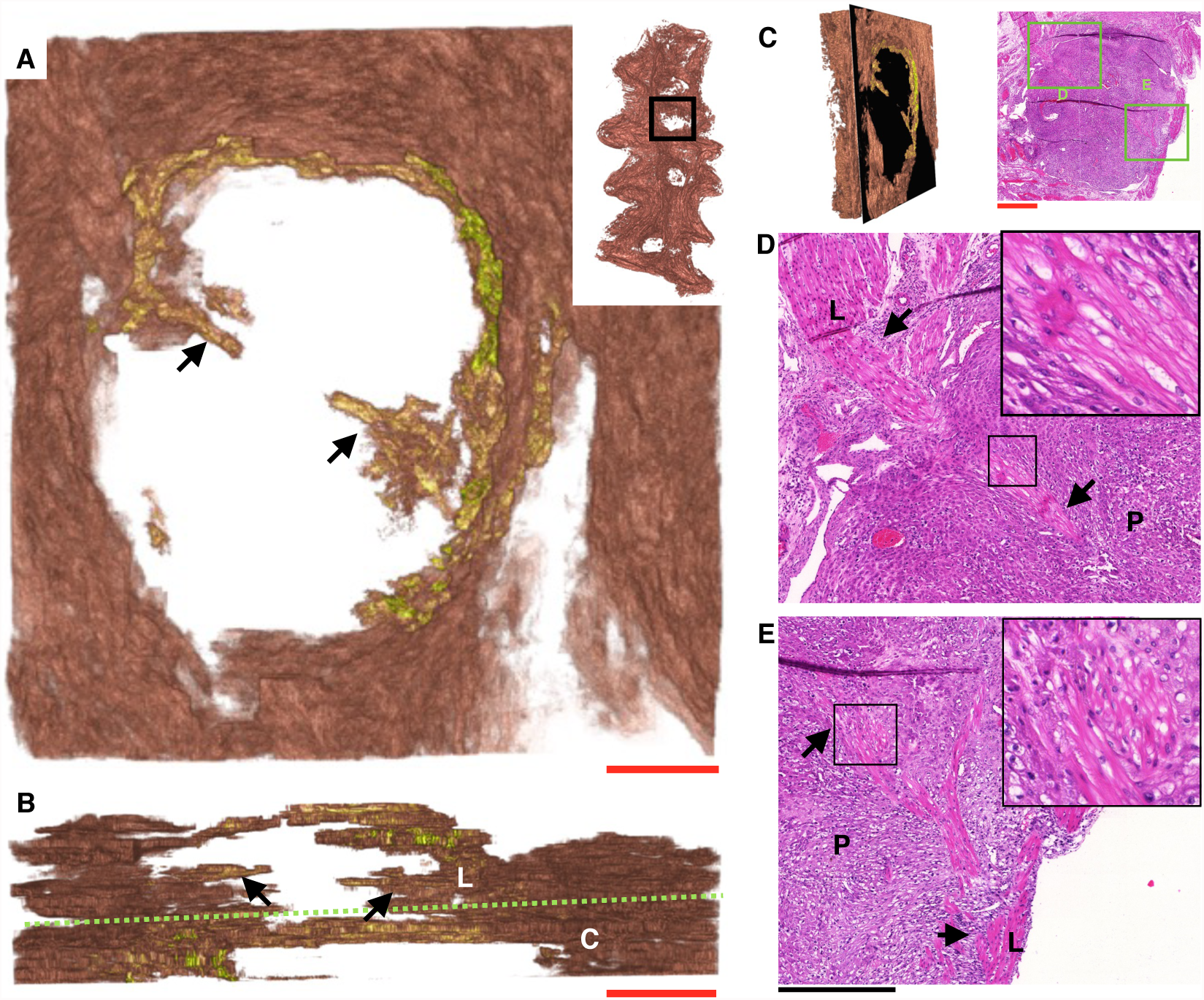
**A:** Three-dimensional reconstruction of myometrial bundles (arrows) present in the placental bed, observed from the serosal side of the tissue. **B:** The same bundles (arrows) viewed from the cervical end of the tissue, with the approximate boundaries between the longitudinal (L) and circular (C) layers indicated by a green dashed line, showing that the bundles are attached to the longitudinal myometrium. **C:** Comparison of reconstructed tissue (**A** & **B**) and histological slides (**D**&**E**). **D** & **E**: histology of bundles shown in **A** & **B**, with the points of attachment to the longitudinal myometrium and farthest extension into the placenta indicated by arrows. Insets show more detail of the boxes indicated, in each case revealing that the cells in these structures have a stellate morphology. The initiation points for electrical activity in this uterus occur consistently in close proximity to the structure shown in **D**. All images (excluding **B**) are oriented with the ovarian end of the tissue at the top of the image. L: longitudinal myometrium, C: circular myometrium, P: placenta. Red scale bars represent 1 mm, black scale bars represent 500 *μ*m.

**Figure 5.**
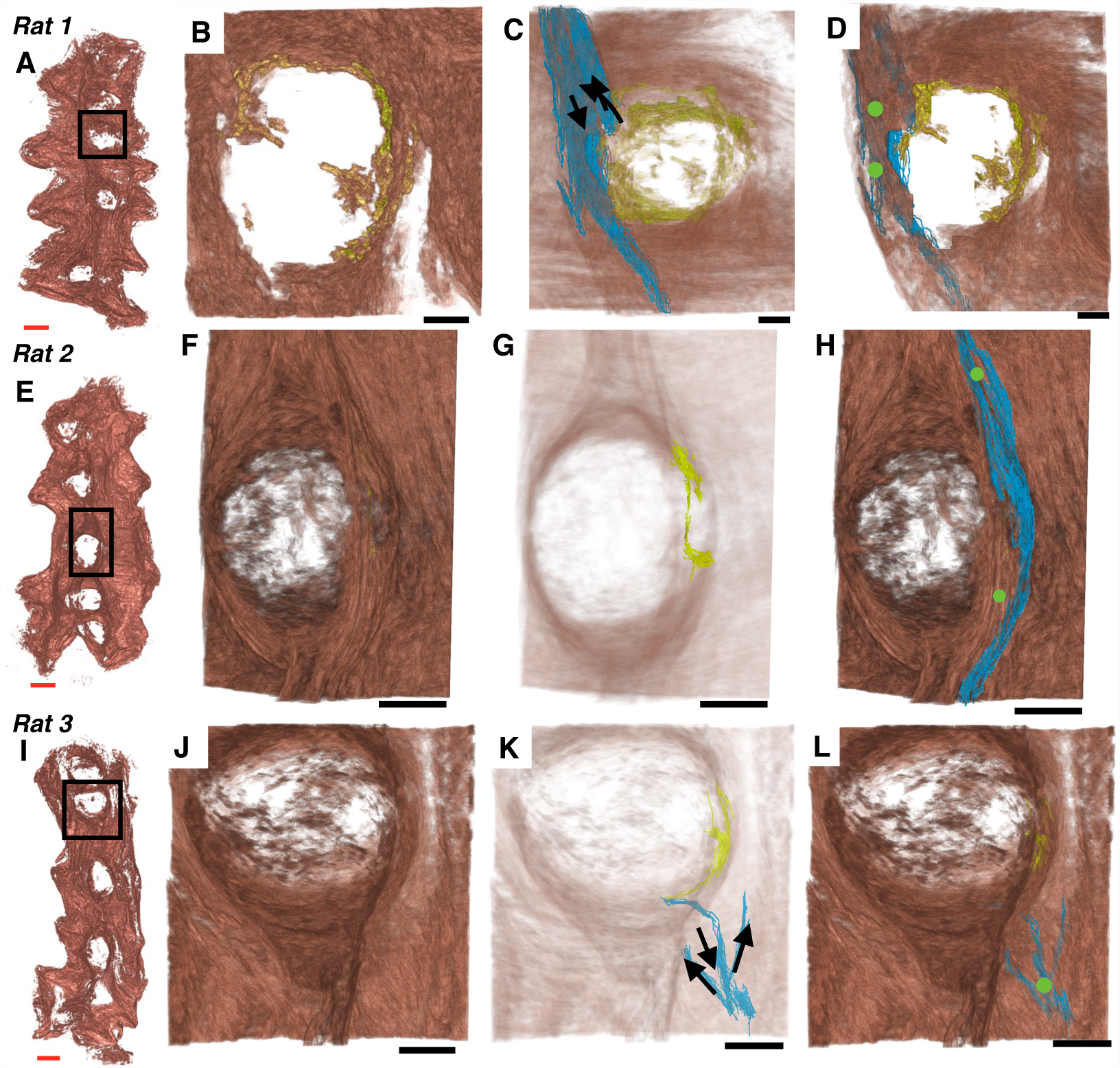
MPPZs associated with excitation activity in relation to the surrounding reconstructed tissue. **A**, **E**, & **I:** Location of the placenta containing the MPPZ. **B**, **F**, & **J**: High-resolution reconstruction of the myometrium adjacent to the placenta containing the MPPZ. The MPPZ is shown in **B**, having been identified in the reconstruction algorithm (see also Fig 4). **C**, **G**, & **K:** MPPZ (yellow) with reconstructed tissue shown at reduced opacity for clarity. The proposed path of excitation (blue) is shown in **C** & **K** with the proposed direction of excitation (arrows). The proposed path is omitted from **G** in order to show the MPPZ clearly. **D**, **H**, & **L:** MPPZ (yellow), proposed path of excitation (blue), and the location of electrodes identified as initiation points in the recordings (green circles). Longitudinal myometrium was present superficial to the MPPZs; it was therefore necessary to identify the path that an excitation wave follows from the MPPZ to the serosal surface, where the electrodes were situated. The proposed paths of excitation were obtained by visual inspection of the histological slides, using the 3-dimensional reconstructions to identify corresponding bundles between slides. The proposed path in the third sample (**K** & **L**) is restricted to the bundle connecting the MPPZ to the myometrium and the shortest unbroken path from this bundle to two adjacent electrodes; adjacent bundles are connected to these bundles, but have been omitted for clarity. Red scale bar represent 5 mm, black scale bars represent 2 mm.

## Discussion

Parturition in the rodent is initiated by a combination of endocrine and paracrine signals. Central to the onset of labor is the systemic withdrawal of progesterone caused by prostanoid-induced collapse of the corpus luteum [2, 21, 41]. The fall in circulating progesterone levels, combined with increased local progesterone resistance [10, 39], activates genes that encode for contraction associated proteins, such as connexin 43 [17] and oxytocin receptor [13,14], that render the myometrium both responsive to excitation stimuli and synchronize contractions. Concomitant with alterations in susceptibility to stimulation, decidual senescence [7,23] and fetal signals [9] increase prostanoid synthesis promoting electrical excitation, voltage-gated calcium entry, and contraction [36] but the precise site of activation has, until now, remained unknown.

In this study, we combined multi-electrode recordings with high-resolution anatomical reconstruction to demonstrate that electrical potentials predominantly originate at specialised interfaces of the myometrium and placenta. Moreover, our results suggest that these myometrial-placental pacemaker zones (MPPZs) form a conduit between each implanted feto-placental unit and the broader myometrial smooth muscle network.

The spatial organisation of the bundles of myometrial smooth muscle in the area of vascular and connective tissue between the longitudinal and circular layers appears to be of particular significance. Bridge-like structures that apparently occur randomly across the tissue, but always in close proximity to the vasculature, form connections between the inner circular myometrium and the outer longitudinal myometrium. The mesometrial-antimesometrial axis governs orientation in rodents; implantation into the anti-mesometrial lumen on day 4 after formation of the vaginal plug orientates the blastocyst for placentation into the mesometrial border [11]. The asymmetry of implantation is encoded by a Wnt5a gradient across the uterine lumen that creates a timed evagination and subsequent implantation crypt [8]. The regular spacing of the implantation crypts is directed by planar cell polarity signalling, as mice deficient in the non-canonical Wnt intermediary *Vangl2* exhibit defective crypt formation and severely compromised pregnancy outcomes [46]. Our data suggests that the asymmetry of patterning, combined with the association of bridges with vasculature, promotes co-localisation of implantation with the vascular supply of the mesometrial axis, and the bridge-like structures of myometrium that provide electrical access to the entire myometrial network. It remains unclear whether the bridge-like structures are formed postnatally concurrently with the circular and longitudinal layers [4] or at the same time as the crypt structures of the lumen [8,11,46]. However, as the vascular network forms before the circular and longitudinal layers [4] we surmise that bridges form postnatally through the paths created by the vascular bed. Our computational reconstructions of the MPPZ bundles suggest that they originate from the invading trophoblast gaining access to, and surrounding, the bridge-like structures of the interconnecting layer. This implies that poor implantation, or implantation outside of the mesometrial axis (*i.e.* not associated with the vascular bed), would lead to poor pregnancy outcomes, as has been demonstrated experimentally in several genetic mouse models [40, 42, 45]. The anatomical restructuring of the myometrium and the formation of the MPPZs described in this study precedes the gestation-dependent alteration in electrical excitability that is conserved across all mammals [6,35]. These changes are mediated by the expression of different classes of potassium channel [3], some of which control resting membrane potential [32] whilst others modulate the action potential waveform to allow for more forceful contractions of longer duration [1, 20, 25, 26, 34, 37]. Thus, the timely activation of the MPPZ at each feto-maternal interface, in a coupled and excitable electrical network, provides an elegant solution for driving contraction from the implantation site towards the cervix to facilitate delivery of multiple fetuses in a polytocus uterus. In conclusion, we have shown that MPPZs represents a thus-far un-recognized structure essential for the coordinated initiation of contractile activity in the rat myometrium.

## Materials and Methods

### Experimental preparation

Virgin Wistar rats (n=3) were time-mated, and pregnancy dated as day 0 of gestation if the sperm cells were observed in the vaginal lavage the next morning. Rats were euthanized (D19-20) by graded CO_2_ inhalation, and the uterine horns were rapidly excised via a midline incision of the abdomen (institutional ethical approval: AE/03/30). The uterine horns were opened longitudinally along the mesometrial border and embryos removed and pinned with the serosal side facing upwards, to a final dimension of approximately 20 mm×50 mm. The tissue was then perfused in modified Tyrode’s solution (mmole/L: 130 NaCl, 4.5 KCl, 2.2 CaCl_2_, 0.6 MgCl_2_, 24.2 NaHCO_3_, 1.2 NaH2PO_4_ and 11 glucose, saturated with carbogen (95% O_2_/5% CO_2_) pH 7.35 ± 0.05, 37 ± 0.5° C) at a rate of 100 ml/min.

### Electrophysiology

The electrophysiological experiments were performed as described in detail by Lammers *et al.* [28]. Electrical recordings were made using a custom rectangular 240-electrodes array (24× 10; 2 mm inter-electrode distance), which covered the entire preparation. The electrodes consisted of Teflon-coated silver wires (0.3 mm diameter, Cooner Wire). Unipolar electrograms were recorded from each individual electrode, with a silver plate located in the tissue bath serving as the common reference electrode. All electrodes were connected through shielded wires to 240 AC amplifiers where the signals were amplified (4000×), filtered (bandwidth 2–400 Hz), digitized (8 bits, 1 KHz sampling rate) and stored on a PC. Recordings were performed for 30 consecutive minutes. After the experiments, signals were digitally filtered (using a 20-point moving average) and displayed on screen. The beginning of every burst was located in time and the location of the first electrical signal in every burst noted.

### Processing multi-electrode array electrograms

The first step is to the determine the characteristics of electrical activity in the tissue from the recordings at each electrode. For each electrode, activity was recorded as a time sequence of voltage values, which is referred to in the following as an *electrogram.* The digitisation described above fixed the time difference between subsequent values as 1 ms. Accordingly, the following shall assume that time is integer-valued, with each time-step equivalent to 1 ms.

An action potential appears in the electrogram as a rapid deflection in the negative direction immediately followed by a rapid deflection in the positive direction, creating a trough-peak pair, as illustrated in Figure 6A [24]. For this reason, it was deemed appropriate to use the difference between a trough and the subsequent peak, known as the peak-to-peak difference [19], to identify action potentials. The peak-to-peak differences were subjected to a thresholding procedure to obtain the timing of each action potential. These action potentials were grouped together to determine the location in time of activity bursts for each electrogram.

**Figure 6.**
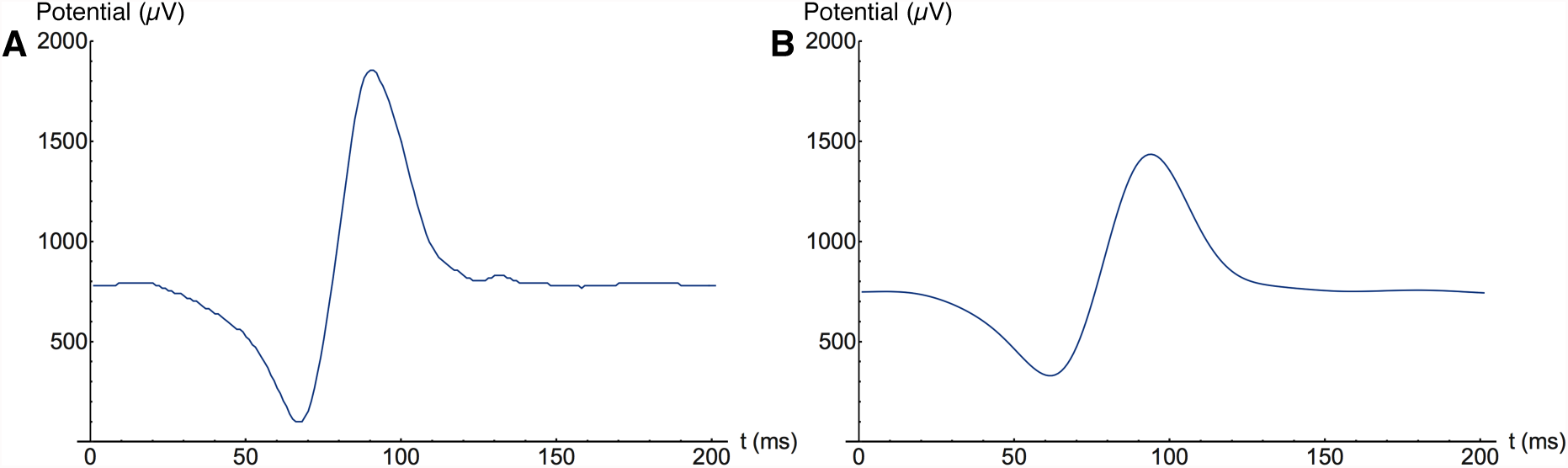
An example of an action potential in an electrogram, both before (**A**) and after (**B**) smoothing.

### Obtaining peak-to-peak differences

In order to reduce noise, a Gaussian filter was applied to the electrogram [5]. The filter used to smooth the data had standard deviation 10 ms and radius 20 ms; these values ensured local smoothing of the electrogram while preserving the general shape of the curves. An example of this smoothing is shown in Figure 6**B**. Although the smoothing can displace the troughs and peaks slightly, this movement is invariably within the bounds of the filter radius, which is sufficiently small to be contained within the width of a peak or trough representing an action potential. Applying this filter to the electrogram yielded a time series of smoothed values which will be denoted by *g*(*t*).

The peaks and troughs of a smoothed electrogram were located by numerically approximating the first derivative. Numerical differentiation is inherently unstable; to control this instability, a seven-point derivative formula was used [5]. The application of this filter to *g*(*t*) produced the estimated derivative *d*(*t*). A time *t* with derivative *d*(*t*) was defined to be a turning point if *d*(*t* – 1) ≠ 0 and

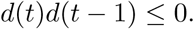

Each turning point was categorised as being either a peak or a trough. To determine whether the turning point was a peak or trough, the second derivative was used. The second derivative was approximated by applying the derivative filter a second time. While it is possible to use a second derivative filter to obtain a more precise estimation of this value [5], a high level of precision is not necessary here and accordingly the second application of the first-derivative filter was used for convenience. Each turning point was categorised as being a trough if the second derivative was less than zero, otherwise it was categorised as being a peak. The Gaussian smoothing, combined with the sharp turning points associated with peaks and troughs from action potentials, ensure that the second derivative is of a sufficiently high magnitude at these points to prevent the error in the derivative estimation from affecting this categorisation. This categorisation includes points of inflection in the peaks. This distinction does not affect the ensuing analysis, because points of inflection are not relevant to action potentials and therefore are generally only present at points which are not part of an action potential.

Each action potential is characterised by a trough followed by a peak. Accordingly, each trough was paired with the next peak to be detected in the time sequence to obtain all possible times at which an action potential may have occurred. A sequence {*p_i_* = *p*(*t_i_*)} of peak-to-peak differences was thus generated, where each *t_i_* is the time at which a trough occurred, and

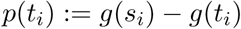

where *s_i_* is the time of the peak paired to the trough at *t_i_.* The effect of this pairing is shown in Figure 7. A thresholding algorithm was applied to this sequence to obtain the action potential times, as described in the following section.

**Figure 7.**
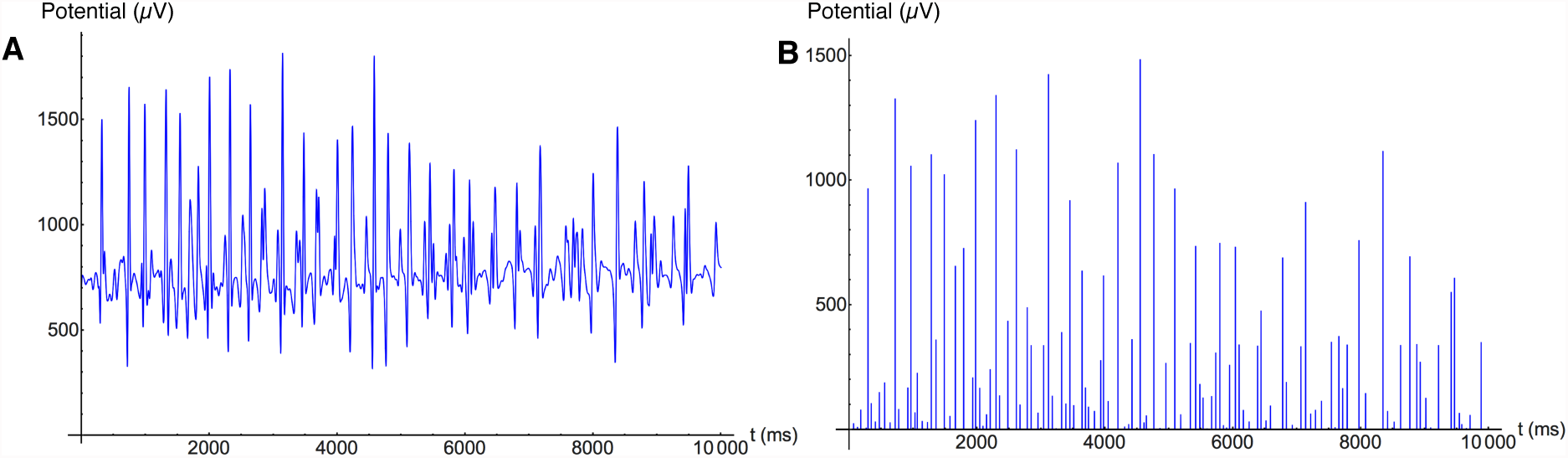
The peak-to-peak difference used to identify action potentials. **A:** A 10-second interval of an electrogram displaying bursting behaviour. **B:** The peak-to-peak differences, with each value placed at the time of the trough’s minimum.

### Identifying action potentials

The peak-to-peak differences that represented action potentials varied in amplitude between electrodes, and therefore the threshold values used to detect the action potential had to be selected separately for each electrode to reflect this amplitude variation. Furthermore, action potential amplitude varied with time in an electrogram, which necessitated the introduction of time-dependent threshold values.

Two forms of thresholding on the peak-to-peak differences were used to determine the action potential times. Both thresholds varied over the duration of the electrogram, with the threshold determined at each point by considering the data within a fixed time interval around the point. The first threshold was a coarse threshold with a large associated time interval to remove low level noise from the peak-to-peak differences, as illustrated in Figure 8**A**. The second threshold refined the set of action potentials by using a localised form of Otsu thresholding [33] with a smaller associated time interval, which removed excess high-amplitude peak-to-peak differences detected over the duration of an action potential, as illustrated in Figure 8**B**.

**Figure 8.**
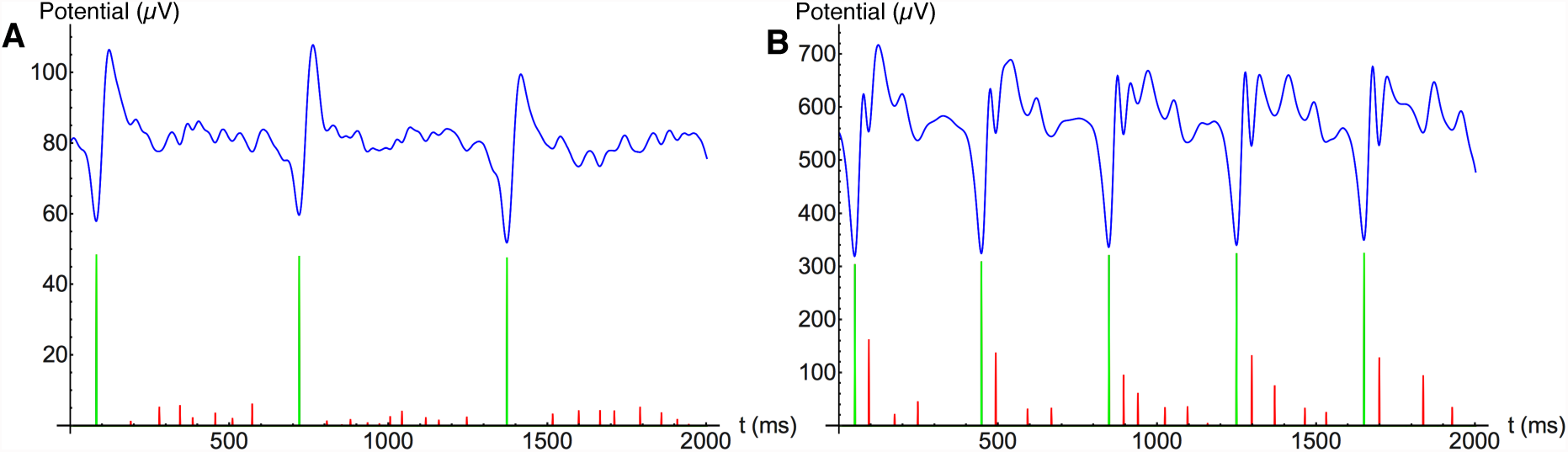
The identification of action potentials from peak-to-peak differences using localised thresholding. Blue represents the smoothed electrogram, red represents peak-to-peak differences, and green represents the peak-to-peak differences greater than the threshold. **A:** The application of a coarse threshold to the peak-to-peak differences removes low-amplitude noise, while maintaining high-amplitude differences. **B:** Action potentials frequently contained multiple high-amplitude peak-to-peak differences, and therefore a refined threshold, based on Otsu thresholding, was required to select a single peak-to-peak difference to represent the action potential.

The coarse threshold was determined by computing the interquartile range of portions of the smoothed electrogram. The aim is to identify extreme peak-to-peak differences, which correspond to action potentials. The set *S*(*t*) of times used to calculate the interquartile range for a time *t* is given by

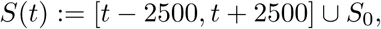

where *S*_0_ is a 5000 ms time interval containing no excitation activity. The inclusion of *S*_0_ in this set provides a baseline for the level of noise, and prevents the threshold from becoming too large in time periods with action potentials of varying amplitudes. The baseline set *S*_0_ was determined for all electrodes by inspecting the beginning and ends of the electrograms: for recordings where all excitation activity occurred after the first 5000 ms at all electrodes, *S*_0_ was set to [0, 5000]. For recordings where excitation activity was observed prior to 5000 ms, *S*_0_ was set to the last 5000 ms of the recording, which contained no activity in all such cases. The threshold at *t* was set to be three times the interquartile range of *S*(*t*). For electrograms with no activity (Figure 9) this threshold was roughly constant and removed the majority of spurious peak-to-peak differences. The threshold varied in electrograms containing bursting activity (Figure 10), and removes small peak-to-peak differences, which correspond to noise, while maintaining the larger differences, which correspond to action potentials.

**Figure 9.**
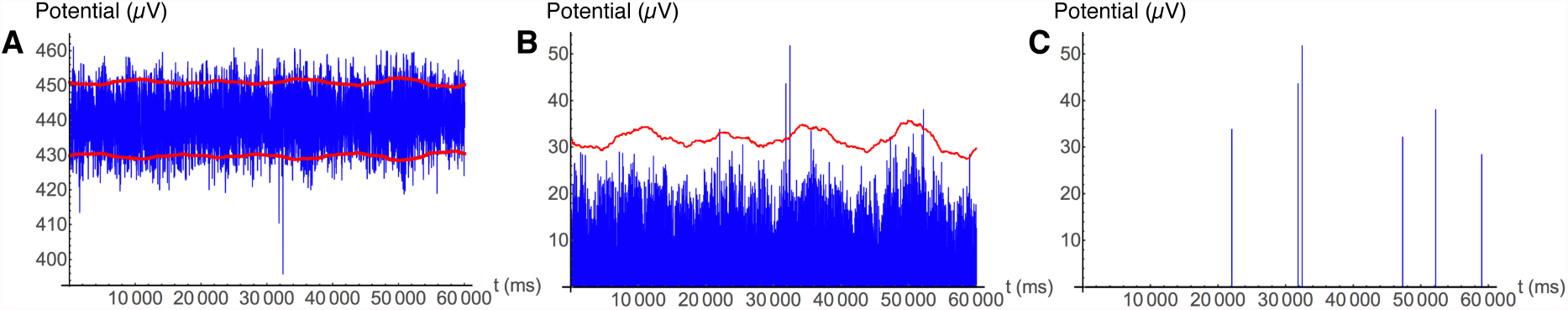
The effect of applying the coarse threshold to an electrogram with no bursting behaviour. **A:** An electrogram with no bursting behaviour, with the first and fourth quartiles of *S*(*t*) marked in red (see text for details). **B:** The peak-to-peak differences, with the threshold equal to three times the interquartile range of *S*(*t*) marked in red. The threshold remains within a range of ~10 *μ*V, which reflects the homogeneity of the input signal. **C:** The result of applying the threshold to **B.** While a few peaks remain, they are not sufficient in number to constitute bursting behaviour, and will therefore be discarded when the bursting behaviour is analysed in a later step.

**Figure 10.**
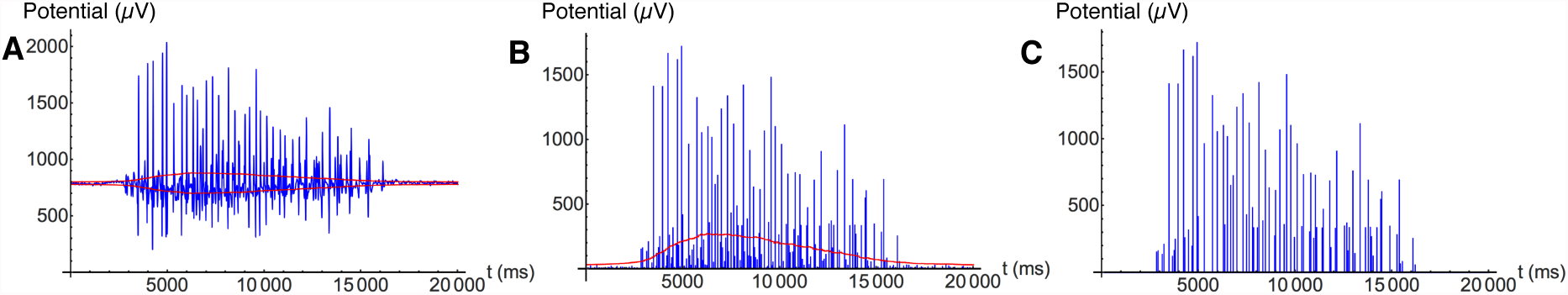
The effect of applying the coarse threshold to an electrogram displaying bursting behaviour. **A:** A 20 s portion of an electrogram showing bursting behaviour, with the first an fourth quartiles of *S*(*t*) marked in red. **B:** The trough-peak pairing, with the threshold equal to three times the interquartile range of *S*(*t*) marked in red. The threshold is increased during a burst, which serves to reduce the level of noise in the output. **C:** The result of applying the threshold to **B.** This threshold removes much of the noise from the bursting activity, but a more refined thresholding technique, based on Otsu thresholding, is required to isolate the action potentials.

The second threshold to be applied to the peak-to-peak differences was determined by applying a localised form of Otsu thresholding [33], which is a general method of determining a threshold that separates foreground from background in image processing [33]. This technique is equally applicable to signal processing, as it does not explicitly rely on spatial data to determine the threshold. The aim of Otsu thresholding is to separate the set *S* = {*p_i_*} into two sets such that the combined spreads of the two sets is minimal. For a threshold value *m*, the sets *S*_0_(*m*) and *S*_1_(*m*) are given by

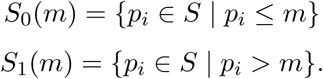

The intra-class variance 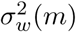 is defined to be

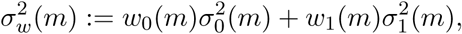

where 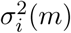 is the variance of *S_i_*(*m*) and *w_i_*(*m*) is the proportion of points belonging to *S_i_*(*m*), given by

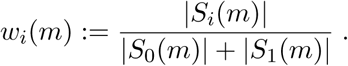

The value of *m* which minimises 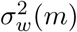 is the point at which the weighted average of the distribution of points in the sets *S_i_*(*m*) is smallest. This minimising value of *m* therefore represents the most likely boundary between two separate distributions of points. Minimising the intra-class variance is equivalent to maximising the inter-class variance 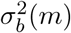 [33], given by

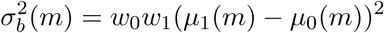

where *μ_i_*(*m*) is the mean of *S_i_*(*m*). This equivalence is apparent from the following relation:

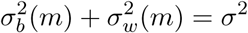

where *σ*^2^ is the variance of *S.* The optimal threshold is the value *m* which minimises *σ_b_*, and this threshold is selected to be the threshold applied to the set *S.*

While this procedure is capable of finding the optimal threshold for the general separation of foreground and background, the problem at hand requires a modified version. The values of {*p_i_*} corresponding to action potentials vary during the burst, as can be seen in Figure 7. In particular, action potentials in one region of the electrogram are not necessarily greater in magnitude than background levels in other regions. For this reason the threshold was set locally, which was achieved in the following way. The time variable was divided into bands of 100 ms, with the threshold calculated separately for each band. For each time band, the set of points used to determine the Otsu threshold was given by points *p_i_* such that *t_i_* was within 200 ms from the time band, as illustrated in Figure 11. This localisation enabled the threshold to be precisely tuned for a short time period, while the extended radius sufficiently increased the sample size to give an accurate threshold value. Using this extended set of values, the threshold in each time band was determined using the Otsu thresholding procedure detailed above.

**Figure 11.**
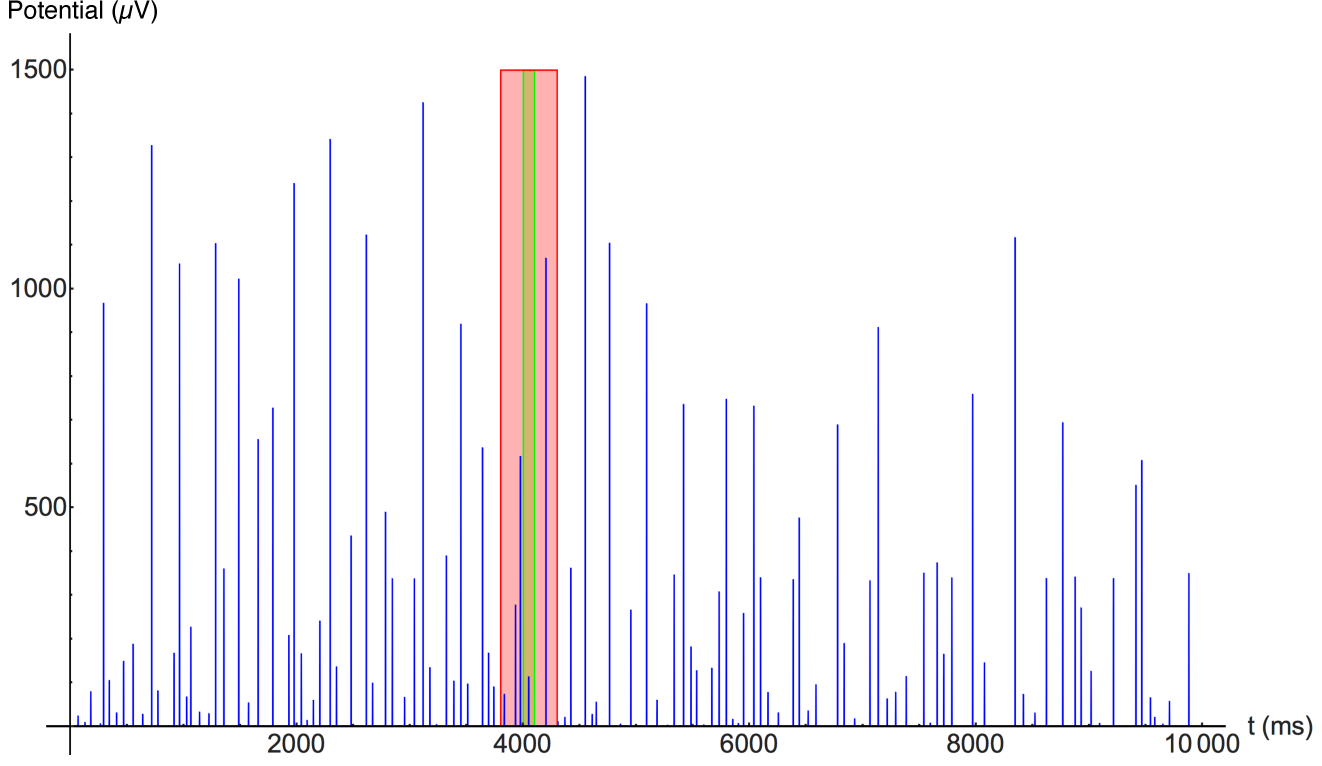
An example of a time interval used to calculate the Otsu threshold of a given time band. The time band marked in green is subject to the Otsu threshold based on values in the interval marked in red.

The modified Otsu thresholding procedure produced a set of localised thresholds as illustrated in Figure 12. This localised thresholding procedure yielded an accurate set of action potential times for each electrogram. These times were used to determine the bursting behaviour of the system as described in the following section.

**Figure 12.**
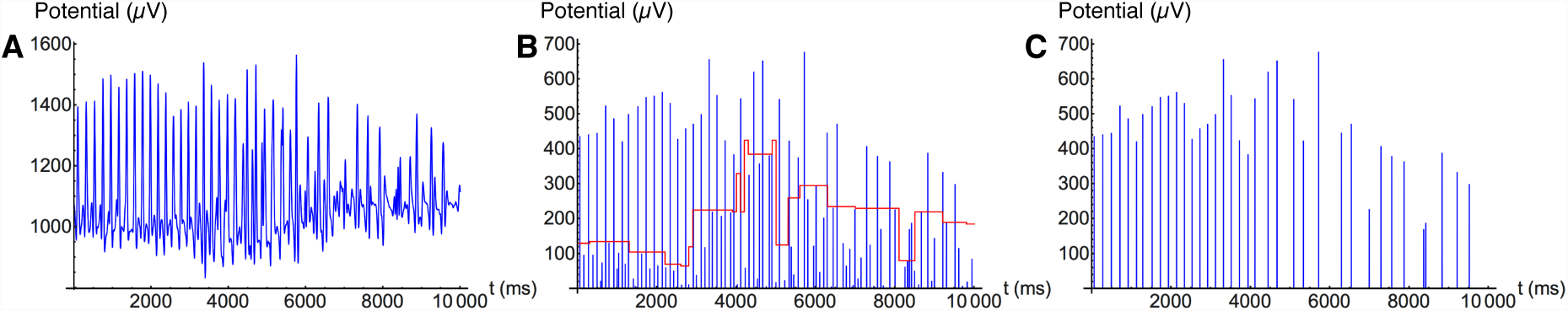
The effect of applying the Otsu threshold to an electrogram displaying bursting behaviour. **A:** A 10-second interval of an electrogram showing bursting behaviour. **B:** The trough-peak pairing, with the Otsu threshold values marked in red. **C:** The result of applying the threshold to **B.** This threshold is adapted to the local peak-to-peak differences, which enables the removal of lower magnitude differences while retaining the differences representing action potentials.

### Determining bursting behaviour

The action potentials detected by the electrode array occur in the form of sustained bursts, as illustrated in Figure 13**A**. A burst can be defined as a set of action potentials occurring in rapid succession [31]. Accordingly, to identify bursting activity from the action potential times a minimum frequency, a maximum time difference, and a minimum time interval were selected. For each electrode with an associated sequence of action potential times 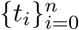, the electrode was said to be recording bursting between the action potential times 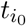 and 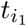 if the subsequence 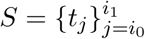 satisfied

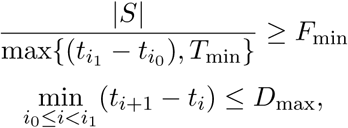

where *T*_min_ is the minimum time interval, *F*_min_ is the minimum frequency, and *D*_max_ is the maximum time difference between action potentials, as listed in Table 1. This definition of bursting behaviour allows multiple bursts to be identified in each electrogram. However, no electrograms showed evidence of multiple burst upon visual inspection. Thus, in the following all electrograms are assumed to contain at most one burst. These criteria were used to identify a burst in the sequence of action potentials in an electrogram, as shown in Figure 13**A**. In some electrograms, a sequence of low-amplitude action potentials within a burst led to the burst being erroneously separated into multiple bursting intervals (Figure 13**B**). In these instances the burst was identified as the subsequence from the start of the first detected burst to the end of the last detected burst.

**Table 1.** Burst detection parameter values.

| Parameter | Value |
| --- | --- |
| $F_{\min}$ | 2 Hz |
| $D_{\max}$ | 1000 ms |
| $T_{\min}$ | 5000 ms |

**Figure 13.**
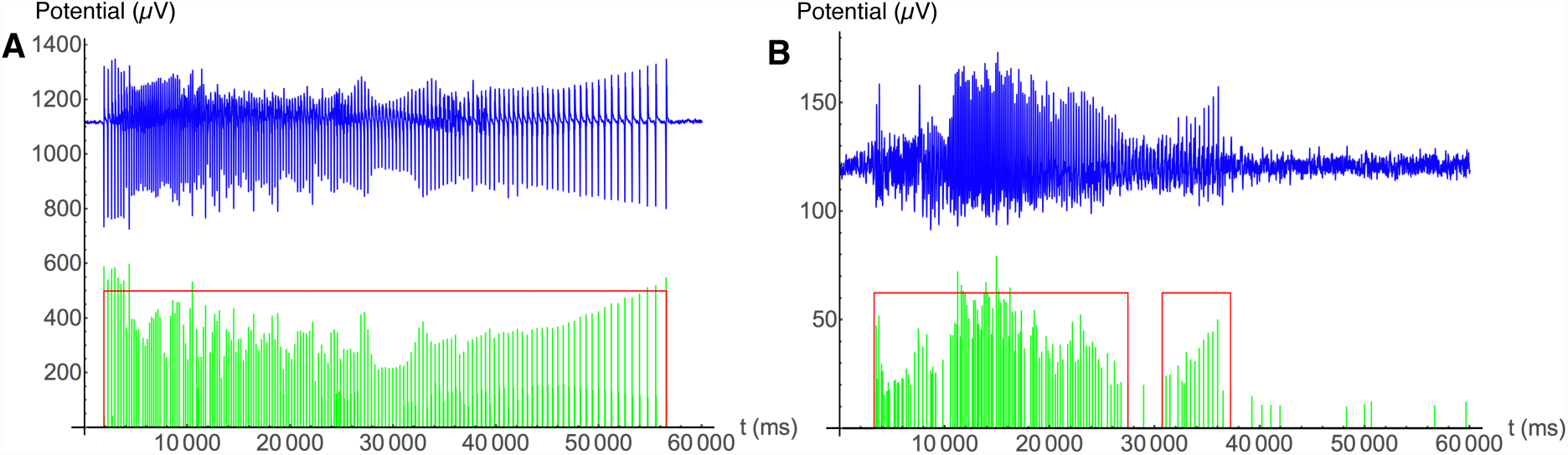
Detection of bursting activity in an electrogram. Blue represents the original electrogram, green represents the peak-to-peak differences of detected action potentials, and red indicates intervals of detected bursting activity. **A:** A burst detected in full as a continuous interval of bursting activity. **B:** A burst partially detected as a set of bursting intervals. This situation occurred when a portion of the electrogram contained action potentials with a reduced amplitude. In this case the complete burst time was taken to be the interval between the first and last times of detected bursting activity.

### Determining propagation behaviour

The bursting activity determined using the process described in the previous section yielded a time sequence of action potentials for each electrode in the multi-electrode array. The next step is to determine how the signal propagates through the tissue on the basis of the bursting data at each electrode. The main challenge is determining which electrodes have recorded bursts from the same propagating wave. To solve this problem, all pairs of electrodes were compared to determine if the bursting behaviour of the given pair is sufficiently similar to be part of the same wave. This requires a suitable quantification of similarity. This pairing of electrodes created a graph of matched electrodes, with each connected component of the graph representing a group of electrodes with a common origin.

It was assumed that if two bursts are part of the same propagating wave, the bursting behaviour in one electrode will be exhibited at a later time point in the other electrode. This similarity may be due to a direct propagation from one electrode to another or from a common origin where the wave has reached one point before the other. The similarity measure, to be detailed below, specifically compared the sequence of action potential times in the respective electrograms. In the following, it is assumed that the excitation waves maintain the same propagation direction for the duration of the recording. For this reason, the comparison measure of electrodes also includes an assumed direction of transmission between the electrodes, and therefore each pair of electrodes was compared in each direction.

The electrogram of an electrode was compared to that of a reference electrode by comparing time intervals of 2000 ms. This time interval was selected to include a sufficient number of action potentials for comparison, while remaining sufficiently short to ensure that any distorting effects of time-dependent changes were minimised. The idea is to slide the action potentials detected in a given time interval backwards in time to compare with the action potentials detected by the reference electrode. The range of times that the interval can be projected back was restricted to the range [50*d*,1000*d*] ms, where *d* is the distance between the electrodes in cm. This range of times represents a propagation speed within the range 1 cm s^−1^ to 20 cm s^−1^, which was selected based on previous observations [38]. Comparison was performed between electrodes that were at most 1 cm apart, to limit the range of projection times and thereby reduce the chance of erroneous matching.

The first step in matching electrodes is to determine if, for a time interval *I* = [*t*_0_, *t*_1_] and projection time *t_P_*, the electrogram of an electrode during *I* resembles the electrogram of a reference electrode during the projected time interval [*t*_0_ − *t_P_*, *t*_1_ − *t_P_*]. Let *α*_0_(*t*) be the time sequence representing action potentials in a reference electrode *e*_0_ such that

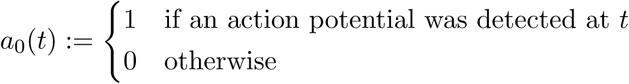

and let *a*_1_ (*t*) be defined similarly for an electrode *e*_1_ being compared to the reference electrode. A time *t* is said to be *matched* for the projection time *t_P_* from *e*_1_ to *e*_0_ if

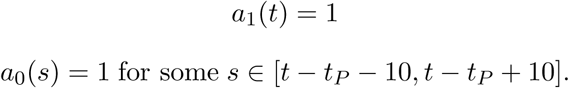

The interval of width 20 ms was selected to allow for any time displacement incurred by the Gaussian smoothing. Similarly, a time *t* is said to be *reverse matched* for the projection time *t_P_* from *e*_1_ to *e*_0_ if

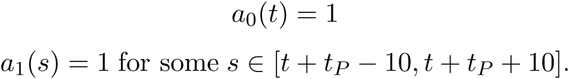

Let *I* = [*t*_0_, *t*_1_] be a time interval containing *N* action potentials in *a*_1_ (*t*), and the interval *I_P_* = [*t*_0_ − *t_P_*, *t*_1_ − *t_P_*] be the projected time interval containing *N_P_* action potentials in *a*_0_(*t*). Then *I* is said to be a *matched* for the projection time *t_P_* from *e*_1_ to *e*_0_ if *N*, *N_P_* ≥ (*t*_1_ − *t*_0_)/500 and

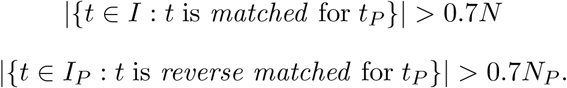

This bound depends on the frequency of action potentials in *I*; action potential frequency is bounded above by approximately 5 Hz, which corresponds to *N* = 10. Thus the above limit permits an interval to be matched with at most 2 action potentials unmatched in each direction at high frequencies (3.5–5 Hz), and at most 1 action potential in each direction at low frequencies (2–3 Hz). For the given range of projection times *R*, an interval *I* is deemed to be a matched interval for an electrode *e*_1_ and reference electrode *e*_0_ if it is matched for some projection *t_P_* ∊ *R* from *e*_1_ to *e*_0_.

This definition of a matched interval was used to determine if an electrode was matched to a reference electrode as follows. The sequence of time intervals {*I_k_*} was defined as follows:

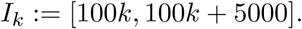

An electrode *e*_1_ was said to be matched to a reference electrode *e*_0_ if the number *N* of matched intervals in this sequence satisfied

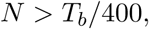

where *T_b_* is the length of the burst detected at the electrode *e*_1_.

This process of matching electrodes produced a graph of the electrodes, where each edge of the graph represents a matched pair of electrodes, as illustrated in Figure 14**A**. These graphs were used to generate isochrone maps, as follows. Each electrode *e_i_* represented in a graph was also identified as exhibiting bursting activity, and therefore has a time *t_i_* at which the burst reached the electrode. These times were coarse-grained into 200 ms bands by assigning an integer value *c_i_* ≥ 0 to each electrode, defined by

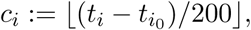

where *t_i0_* is the minimum time *t_i_* and [∗] rounds down to the nearest integer. Each connected component of the graph containing more than two nodes was taken to be a collection of electrodes which exhibit bursting behaviour as part of a common excitation wave. For each connected component of the graph, a closed curve was drawn for each integer *c* > 0 such that

i. all electrodes *e_i_* in the connected component with *c_i_* ≤ *c* are contained within the curve;
ii. all points within 1 mm of an electrode given in (i) are contained within the curve;
iii. the area enclosed by the curve is minimal subject to (i)–(ii).

**Figure 14.**
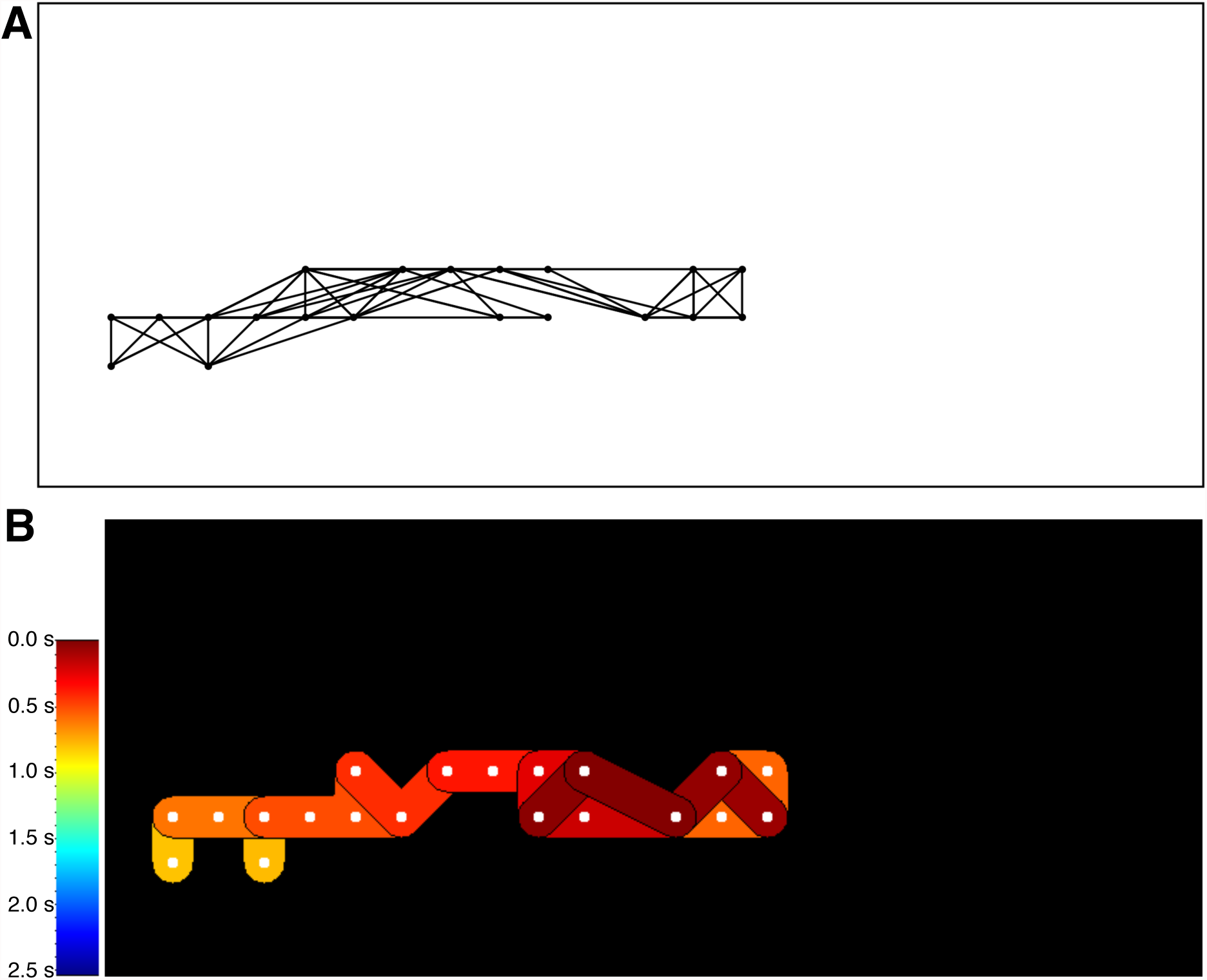
**A:** A graph representing the electrode matching. Each node is positioned at an electrode that has been matched to another electrode, and edges represent these matches. **B:** The isochrone map generated from network shown in **A.** The isochrones are coloured according to the time at which the burst is first detected by the electrode, with red corresponding to earliest and yellow corresponding to latest.

This line is the isochrone for time 200*c* ms. Criterion (iii) serves to prevent the isochrone from including portions of the tissue which are not excited before the given time. The areas enclosed by these isochrones were colour-coded, as shown in Figure 14**B**. These isochrone maps visualise the spread of excitation, and were compared to three-dimensional reconstructions of the tissue to identify structures associated with initiation and termination of the excitation waves.

The methods detailed in this section provide the tools for automated analysis of multi-electrode array recordings. The processing of each electrogram provides a representation of the times at which action potentials occur, and whether the electrogram exhibits bursting behaviour. The analysis of the array as a whole enables the generation of isochrone maps, which show initiation points of propagating bursts, along with locations where propagation is blocked. To ensure accuracy of these isochrone maps, all recordings were visually inspected and the isochrone maps amended where necessary.

### High-resolution reconstruction of myometrial smooth muscle

Two types of structures were identified in histological slides taken from pregnant rat uteri, namely bridge-like structures between the circular and longitudinal layers of the myometrium, referred to as *bridges*, and fibrous structures in the placenta associated with pacemaker activity, referred to as *myometrial-placental pacemaker zones* (MPPZs). In order to study the architecture of these structures at the requisite level of detail, it was necessary to generate a high-resolution three-dimensional reconstruction of the tissue. Methods described in a previous publication [30] constitute an image analysis pipeline for generating a three-dimensional reconstruction of the myometrium from serial sections at a resolution of ~ 50 *μ*m per voxel length. The structures observed in the present paper are generally finer than this, with widths as small as 20 *μ*m, and accordingly precise rendering of such structures requires a higher resolution. The reconstruction techniques described previously [30] specifically identify nuclei in the histological sections. The methods described in the following exploit the orientation of these nuclei to generate a reconstruction of the myometrium at a resolution of 10 *μ*m per voxel length. This reconstruction allows the visualisation of the connectivity between the longitudinal and circular layers and, with some manual adjustment, the MPPZs.

The histological slides used to generate this high-resolution three-dimensional reconstruction were registered and the nuclei identified as previously documented [30]. The slides were obtained by slicing the tissue sample into serial sections 5 *μ*m thick and discarding any slides that were too severely distorted by sectioning to register properly (58%–72% depending on the tissue block), yielding an unevenly-spaced stack of useable slides. In each slide the nuclei were automatically identified and the position, size, and orientation were recorded. These slides were registered to obtain a three-dimensional record of the nuclei in the tissue, with maximal *z*-resolution 5 *μ*m and planar resolution ~ 0.5 *μ*m (here, ‘planar’ refers to the plane of the slides, identified as (*x*, *y*) with *z* being the axis perpendicular to this plane.

### Generating high-resolution reconstructions

The high-resolution reconstruction of myometrial smooth muscle tissue was obtained by identifying which nuclei were contained in smooth muscle tissue, as opposed to surrounding tissues, and generating a representation of the fibrous structure based on the position and orientation of these nuclei. In particular, the algorithm was designed to differentiate between smooth muscle nuclei and nuclei contained in the placental and connective tissues. These tissue types were filtered out because they tend to be present in close proximity to the features to be identified; placental tissue surrounds the MPPZs, while bridges between circular and longitudinal myometrium are located in the connective tissue between the layers. Vasculature is also present between the layers of the myometrium [4] but was not filtered out of the reconstruction, which motivated an additional precautionary step in which any bridges between the layers observed in the reconstruction were also identified in the histological slides to rule out the possibility that they are in fact vascular structures. The filtering of the nuclei was achieved by considering the local homogeneity of nuclear direction and nuclear density in the area surrounding each nucleus; local nuclear direction is more homogeneous in smooth muscle than in the other tissue types, while the shape of the smooth muscle cells imposes constraints on the local density, as shown in Figure 15.

**Figure 15.**
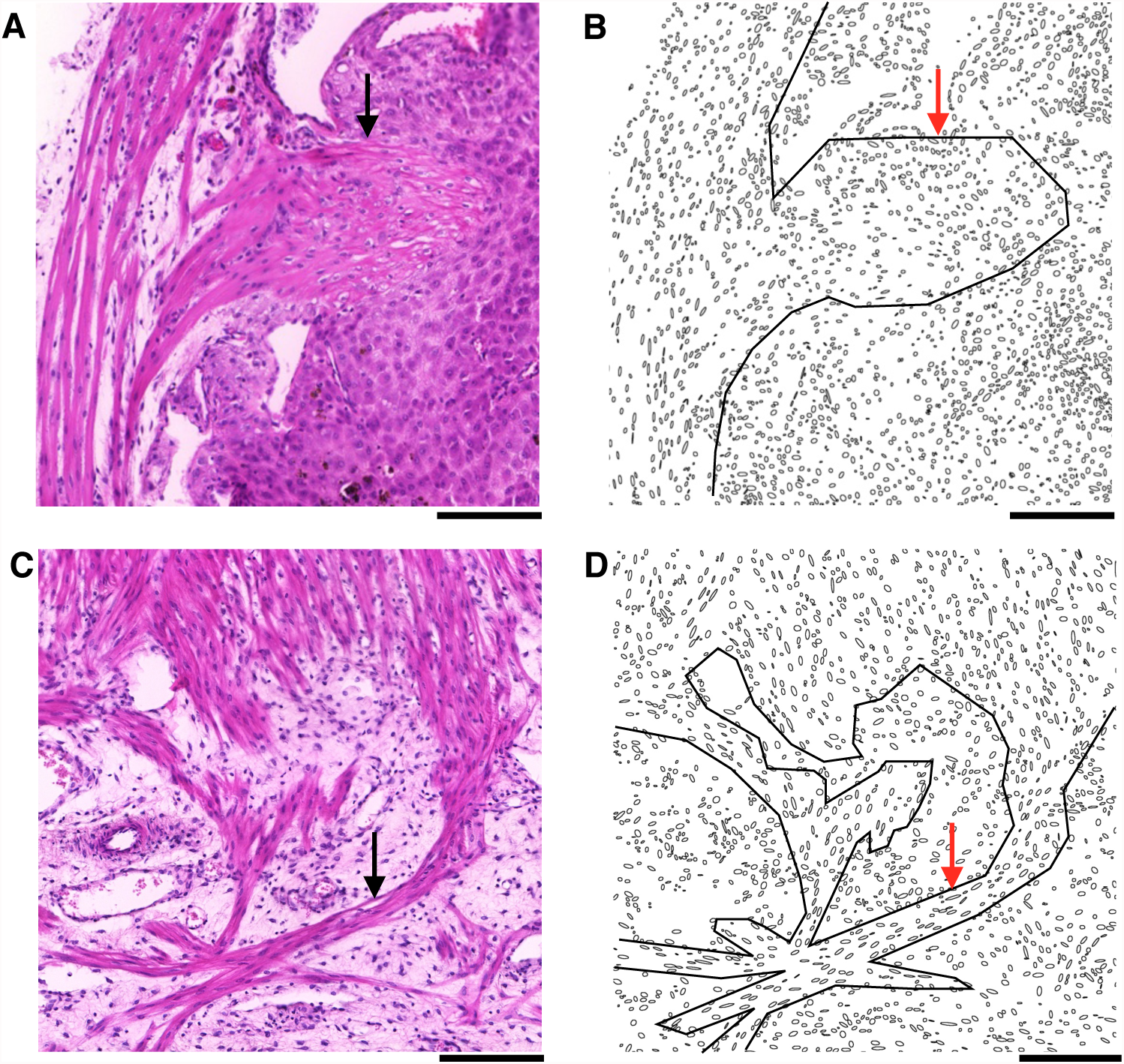
Nuclei in smooth muscle, placental, and connective tissue. The orientation of nuclei in smooth muscle tissue is locally homogeneous, which is generally not the case in placental **(A** & **B)** and connective tissue **(C** & **D).** This local homogeneity is also present in MPPZs (Arrows in **A** & **B)** and bridges (Arrows in **C** & **D).** Additionally, nuclear density is more variable in the placenta and connective tissue than in smooth muscle tissue. Black lines in **B** & **D** represent the approximate boundary of the smooth muscle tissue. **A:** Histological image of smooth muscle and placental tissue. **B:** Nuclei present in **A. C:** Histological image of smooth muscle and connective tissue. **D:** Nuclei present in **C.** Scale bars represent 200 *μ*m.

The procedure for generating a high-resolution reconstruction from the input stack is as follows. Each slide was mapped onto an image representing the nuclear direction, with each pixel representing a 10 *μ*m × 10 *μ*m area of the slide. These angle images were filtered to remove nuclei that did not satisfy the density and homogeneity criteria for smooth muscle tissue. Each of these remaining nuclei was assigned a *score* (defined below), and the fibrous structure was marked in each slide based on the position, orientation, and score of each nucleus, yielding a visual representation of the smooth muscle tissue. MPPZs were identified in each slide by visual comparison with the histological slides. The fibrous structure and manually-identified MPPZs from each slide were combined to produce a volume representing the tissue, with resolution 10 *μ*m per voxel length.

### Generating angle images

Each slide in the stack was coarse-grained to generate an angle image *A* with pixels corresponding to a grid of 10 *μ*m × 10 *μ*m areas in the slide. The nuclei in the slide were filtered by size, with maximum and minimum nuclear size given in Table 2. Let *A_S_*(*p*) be the area in the slide corresponding to the pixel *p.* If *A_s_*(*p*) contains one or more nuclei, then let *θ*(*p*) ∈ [0,180) be the angle that a randomly selected nucleus in *A_s_*(*p*) makes with the *x*-axis. We define

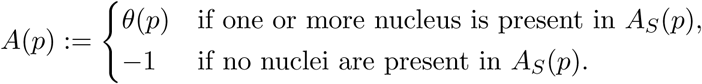

**Table 2.** Parameter values used to generate high-resolution reconstructions.

| Parameter | Value |
| --- | --- |
| Minimum nuclear size | 10.4 $\mu\text{m}^2$ |
| Maximum nuclear size | 62.1 $\mu\text{m}^2$ |
| $d_{\max}$ | 1 pixel lengths |
| $\lambda_{\max}$ | 15 pixel lengths |
| Homogeneity score threshold | 0.90 |
| Maximum density | 0.33 |
| Minimum density | 0.10 |
| $l_r$ | 6 pixel lengths |
| $t_r$ | 1 pixel lengths |
| $z_r$ | 1 pixel lengths |
| Smooth score threshold | 0.05 |

At a resolution of 10 *μ*m × 10 *μ*m, it is unlikely that more than one nucleus is present in any given area *A_S_*(*p*), and therefore the information discarded in the mapping of such areas onto *A* was minimal. Thus, in the following, the cases where multiple nuclei are present in *A_S_*(*p*) are ignored and a pixel *p* is said to correspond to a nucleus if *A*(*p*) ≥ 0. The pixel value of −1 represents empty areas; this value was only used as a marker and not included in any numerical calculations.

### Generating score images

For each pixel *p* corresponding to a nucleus, a rectangle *R_p_* of length 30 pixels (300 *μ*m) and width 3 pixels (30 *μ*m) was drawn along the pixel direction, centred at *p*, as illustrated in Figure 16. More formally, for a line *L_p_*(*λ*) given by

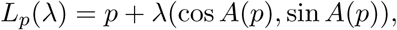

let

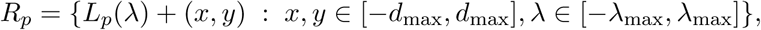

where *d*_max_ and *λ*_max_ are given in Table 2. The homogeneity score *H*(*R_p_*) of the rectangle was calculated as follows:

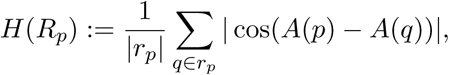

where *r_p_* is the set of all pixels in *R_p_* that correspond to cell nuclei. This quantity is the average scalar product of nuclear direction with the direction at *p*, and represents how well the nuclei in *R_p_* align with the nucleus corresponding to *p.* The density of *R_p_* is given by

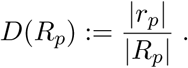

**Figure 16.**
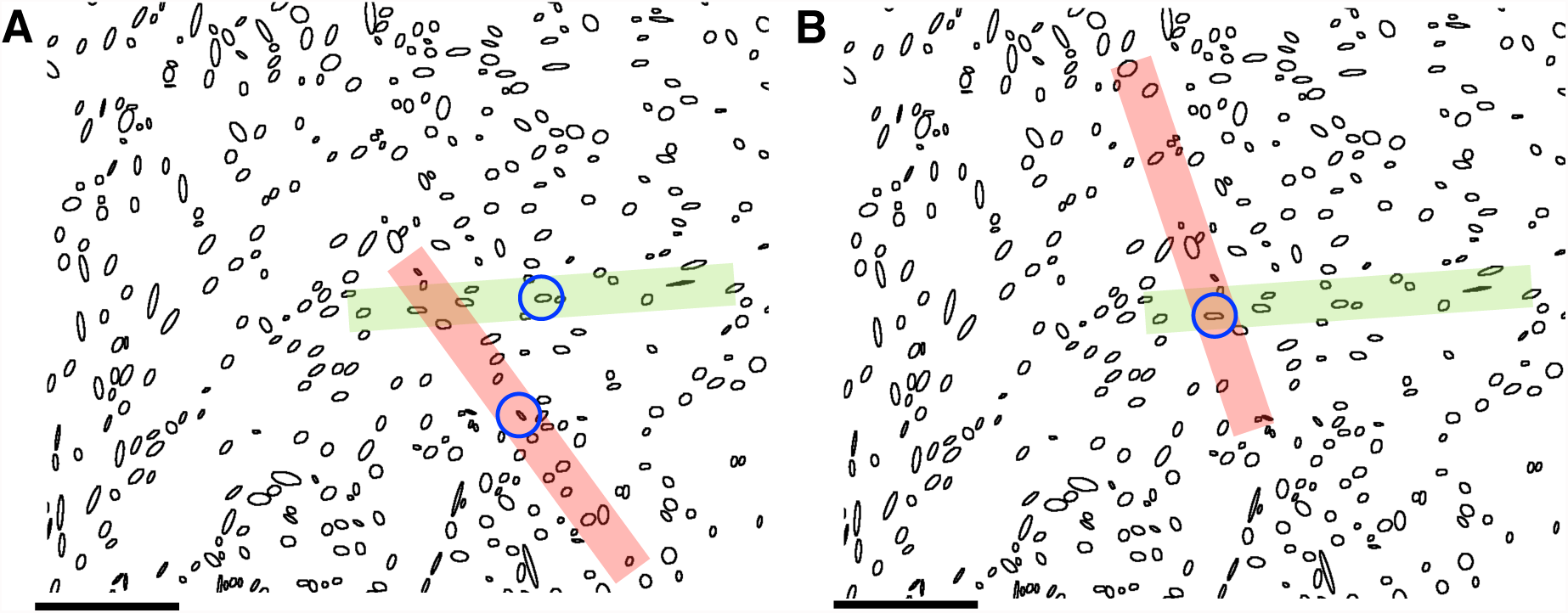
Rectangles used to generate score images. **A:** The initial selection of valid areas. Rectangles are placed over each nucleus along the direction given by the nucleus. The blue rings indicate the two nuclei that correspond to the rectangles shown. The green rectangle contains nuclei that closely approximate the direction of the central nucleus, so this rectangle is categorised as being within smooth muscle tissue. Conversely the red rectangle contains nuclei that show little similarity in terms of orientation to the central nucleus, so this rectangle is not categorised as being within smooth muscle tissue. **B:** Scoring valid nuclei. For a given nucleus, the rectangles that contained this nucleus and were deemed to represent fibrous tissue by the process illustrated in **A** were compared to the nucleus. The rectangle with the closest orientation to the nucleus was selected to generate the score of the nuclei (see text). In this example, the nucleus (blue ring) is closer in orientation to the green rectangle than the red, and thus the green rectangle is used to generate the score.

Threshold values for *H*(*R_p_*) and *D*(*R_p_*) are listed in Table 2. If the value of *H*(*R_p_*) was above the given score threshold and the value of *D*(*R_p_*) was within the given density range, then the rectangle *R_p_* was deemed to represent smooth muscle tissue with bundle direction approximated by the direction at *p*, as illustrated in Figure 16**A**. The set 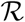 of all such rectangles was used to create a homogeneity score image as follows. Each pixel *q* corresponding to a nucleus and contained in at least one rectangle in 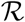 was assigned the score

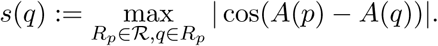

This score represents how well the given nucleus is aligned with a rectangle identified as being aligned with the direction of the local bundle of smooth muscle tissue, and therefore is a measure of how well the nucleus fits into the category of smooth muscle tissue, as shown in Figure 16**B**. This scoring function was used to generate the score image *I*:

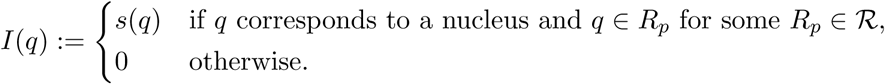

This score image highlights pixels corresponding to nuclei identified as belonging to a smooth muscle bundle, with each pixel assigned a value in [0, 1] representing how closely the orientation of the corresponding nucleus aligns with the local bundle direction. This representation also highlights pixels corresponding to nuclei in MPPZs and bridges: both of these structures have locally homogeneous nuclear direction and similar density to smooth muscle bundles (Figure 15). Additionally the rectangles were constructed such that their width (3 pixels) is similar to the minimum width of these structures (~ 2 pixels), which ensures that a rectangle corresponding to a nucleus in one of these structures is generally contained within the structure.

### Visualising smooth muscle cells

The score images are sparsely populated with positive values because positive values correspond to a nucleus situated in a smooth muscle cell, and smooth muscle cell cross-sections in the slides are typically 5–10 times larger than the area represented by an individual pixel. To visualise bundles, anisotropic Gaussian smoothing was applied [18]. The smoothing kernel *G* was defined as follows:

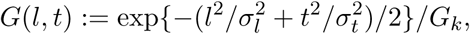

where *σ_l_* is the longitudinal standard deviation of *G* and *σ_t_* is the transverse standard deviation of *G*, as given in in Table 2, *l* ∊ [−*σ_l_*, *σ_l_*] and *t* ∊ [−*σ_t_*, *σ_t_*] are integers, and *G_k_* is a normalising factor:

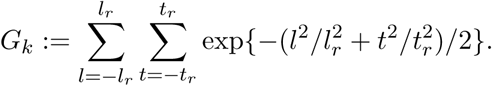

The aim is to rotate apply *G* at each point of a score image *I* such that the longitudinal length of *G* is aligned with the local fiber direction. For a pixel *p* with an assigned angle *θ*, define the image *I_p_* as follows:

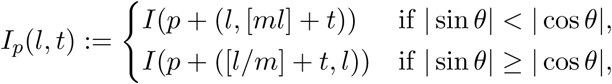

where [·] rounds to the nearest integer, and

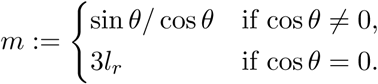

This image is constructed to extend out from *p* at angle *θ* in the plane. The images *I_p_* are only defined for pixels corresponding to nuclei; to extend this definition to the entire image, each pixel not corresponding to a nucleus was assigned the direction of the nearest pixel corresponding to a nucleus. The smoothed image *J* is thus given by

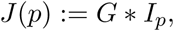

where ∗ represents convolution.

Finally, the stack of images were smoothed in the *z*-direction to improve the reliability of each image. The sequence of smoothed score images {*I_n_*} representing slides in the stack was taken to be uniformly spaced slices through a score volume *V_S_.* This representation somewhat distorted because the slides are not evenly spaced; however, this should not be considered to be a serious problem given that the aim is to reinforce the visualisation for each image by smoothing with images that are adjacent in the stack. A one-dimensional Gaussian filter with radius 1 voxel and standard deviation 1 voxel was applied to *V_S_* in the *z*-direction, where 1 voxel represents the separation of adjacent slides in the stack. The result is the smoothed volume *V.*

### Visual identification of MPPZs

The detection of MPPZs required visual comparison to exclude any placental tissue that has similar density and directionality properties to smooth muscle. For each implantation site, a subvolume containing the implantation site and adjacent smooth muscle tissue of width and height roughly twice that of the implantation site was used as the input for this procedure. The aim is to create a set of possible MPPZs in the placenta and manually identify which of these structures correspond to MPPZs in the histological slides. This method is advantageous over identifying and highlighting each MPPZ manually, because it allows the whole three-dimensional structure of an MPPZ to be identified from visual inspection of just one slide. This serves to reduce the time required to reconstruct the whole MPPZ, while also providing direct correspondence between cross-sectional slices of an MPPZ located in multiple slides.

For a subvolume containing an implantation site, the voxels with positive score were reduced to those connected to the myometrium adjacent to the implantation site. This was achieved by separating voxels into sets *S_i_* of voxels that have score value above the smooth score threshold value given in Table 2, and are connected by adjacency: for any two points *p*, *q*, ∊ *S_i_*, there exists a connected path represented by a sequence {*p*_0_ = *p*, *p*_1_, …, *p_n_* = *q*} ⊂ *S_i_* such that

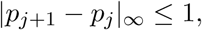

where | · |*_∞_* is the maximal absolute component of a vector, and *V*(*p_j_*) is above the threshold value for all *j.* Provided that the volume bounds around the placenta are sufficiently wide, the largest of these sets 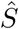 will include the smooth muscle outside the placenta and any potential MPPZs attached to it. Accordingly, all points outside this maximal set were assigned a score of 0. This step serves to eliminate erroneously highlighted placental tissue and aid manual selection of the MPPZs.

An MPPZ is represented in this reconstruction by a cluster of points in the set 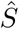 that extends from the myometrium into the placenta. The next step is to segment the points in 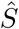 that are situated in the placenta into discrete clusters, so that clusters corresponding to MPPZs can be selected manually based on visual comparison with the histological slides (Figure 17). This visual comparison only requires manual inspection of a sparse selection of slides: an MPPZ intersecting multiple slides only needs to be identified in one slide.

**Figure 17.**
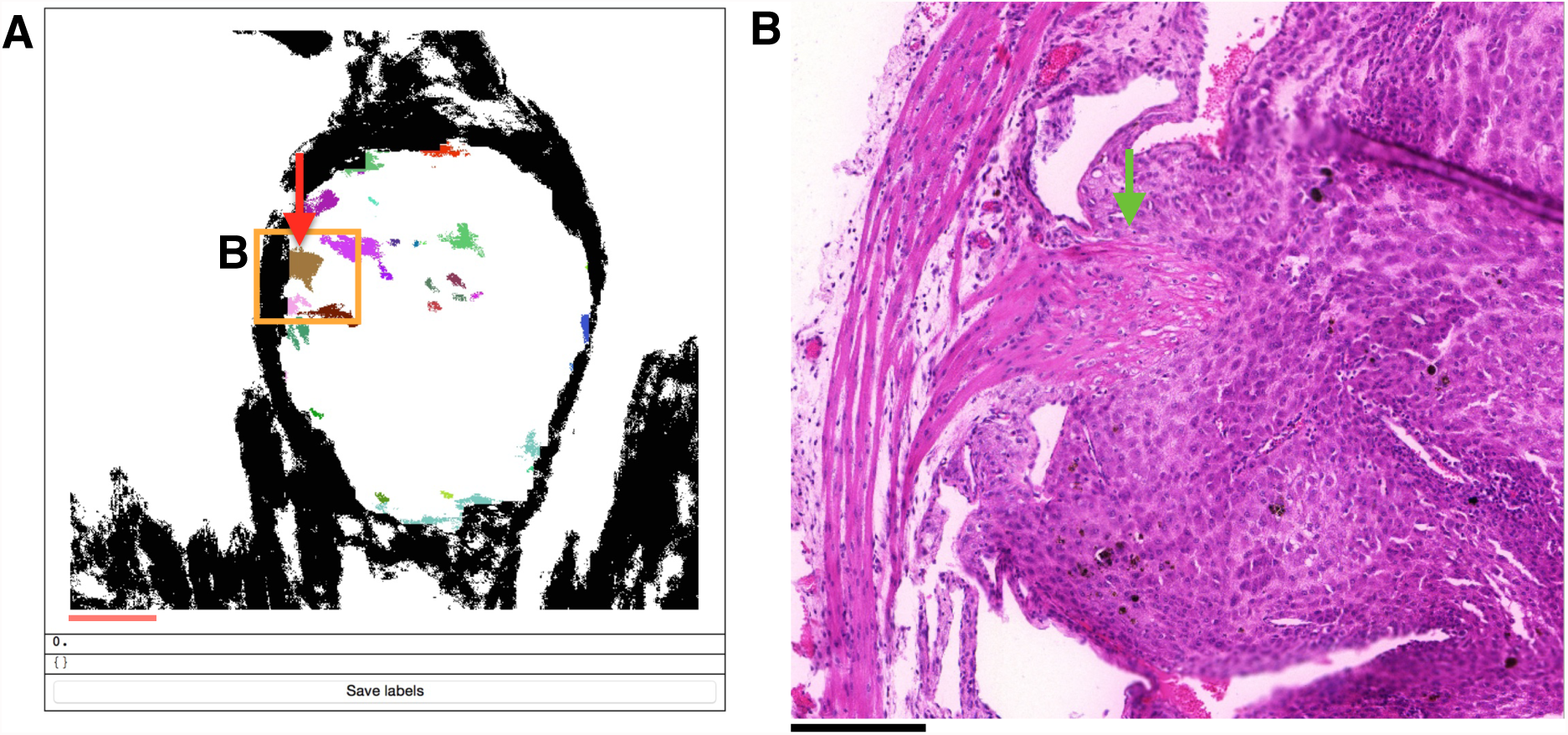
Manual selection of MPPZs. **A:** The user interface used to select MPPZs. The upper panel displays segmented clusters in arbirtrary colours for the given slide; the lower panels are used to write clusters selected by the user to generate the final volume. **B:** A small portion of the implantation site in the given slide. The MPPZ that is indicated in **B** by the green arrow corresponds to the cluster indicated by the red arrow in **A**, and is selected for the final representation. All other clusters in the area corresponding to **B** do not correspond to discrete well-organised structures, as can be seen in the histological image, and thus are not selected. The clusters are three-dimensional: the highlighted cluster is part of a larger structure extending through multiple slides. Therefore, manually identifying this structure in this one slide leads to the automatic selection of the same structure in all other slides. This allows the human operator to verify the MPPZs on a sparse selection of slides and use the clustering to interpolate the overall structure.

The first step in this cluster segmentation is to identify the portion of space in the reconstruction corresponding to placental tissue, which was achieved by identifying the area interior to 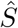 in each slide. For a slide at position *z* given by the set of image points *I_z_* and containing the points 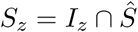, let 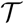 be the index set of connected components of *I_z_* \ *S_z_*: each 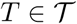 is connected by adjacency in the same manner as for *S_i_* above; for *T*_1_, 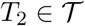 with *T*_1_ ≠ *T*_2_:

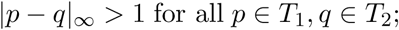

and

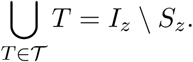

The interior *T_z_* of *S_z_* was defined to be the union of all sets 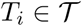 that contained no points at the edge of the image and had size greater than |*I_z_*|/50. In some slides the interior may not be well-defined due to a break in the smooth muscle tissue surrounding the placenta, as shown in Figure 18. In these situations, most or all of the placental tissue will not be present in *T_z_*, and therefore *T_z_* will be reduced in size. Such cases were identified as slides with *|T_z_|* < |*I_z_*|/8. When this criterion was fulfilled, the interior of the nearest slide with a well-defined interior was taken to be the interior of the given slide. The set of all interior points *T* in *V* was taken to be the union of these interior sets:

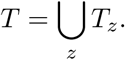

**Figure 18.**
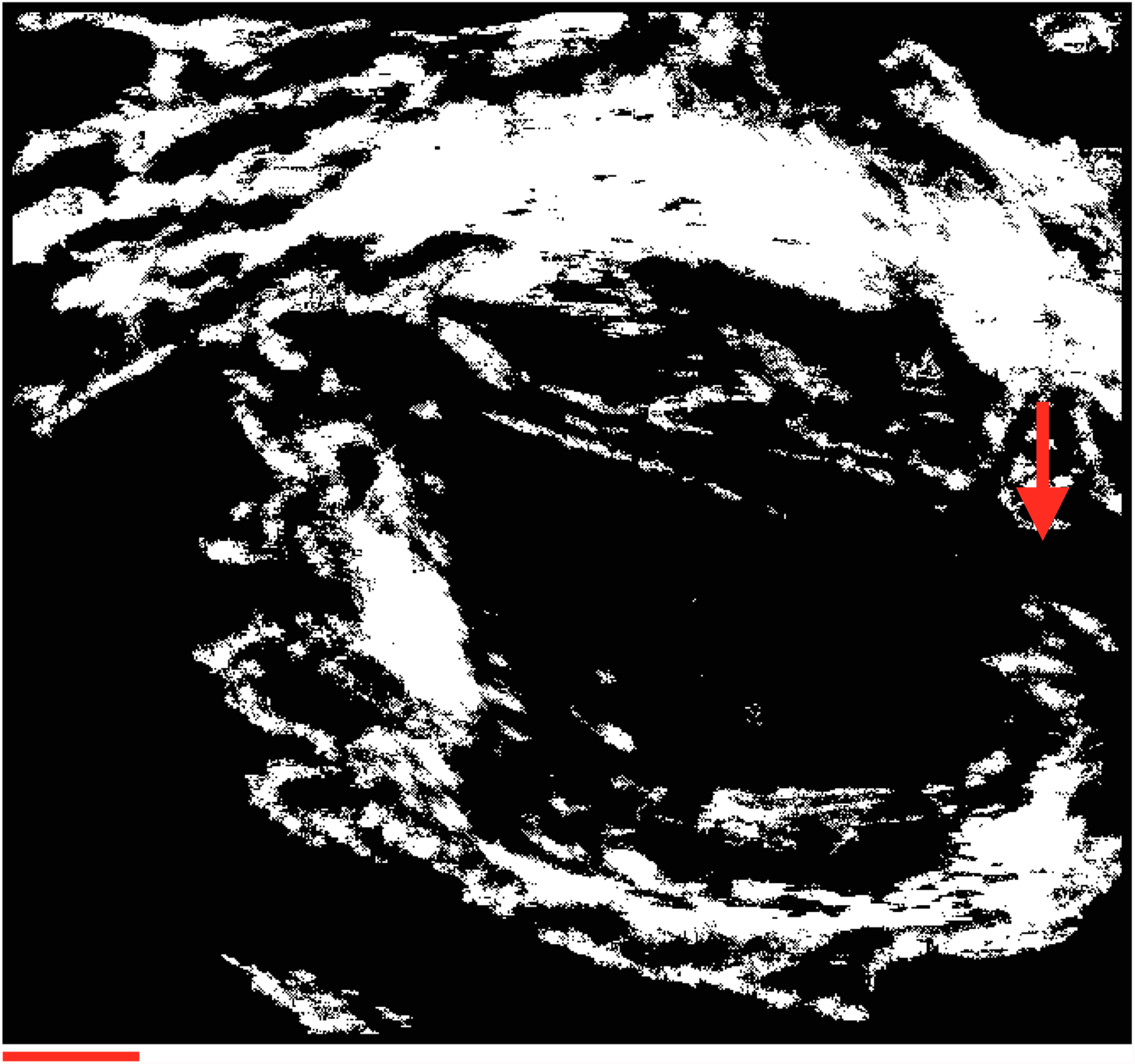
An example where the interior of a structure cannot be identified. The outer structure does not completely enclose the interior, with a gap in the smooth muscle indicated by a red arrow. This means that the interior is not well-defined in this image.

The points in 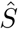 that were contained within the placenta were found by dilating and eroding the interior volume [43]. The set *T* was dilated to yield *T*′:

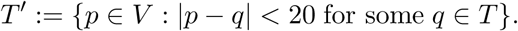

This set was subsequently eroded to yield *T*″:

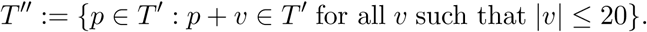

This corresponds to expanding and contracting the interior volume by ~ 200 *μ*m. Any cluster in 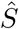 extending into the interior volume of width less than ~ 400 *μ*m would be incorporated into *T*′ upon dilation, but remain in *T*″ after erosion. Thus, the points in 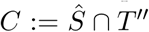 were taken to be the set of points corresponding to possible MPPZs.

Clusters of points in *C* were segmented using hysteresis thresholding: for any point *p* ∊ *C* with value above the upper threshold *t*_upper_, the cluster *C_p_* is the connected set in *C* with values above the lower threshold *t*_lower_. These clusters represent possible MPPZs, and were visually compared to the histological slides as follows. For a slide to be inspected, the cross-sections of all clusters intersecting this slide were displayed in a *Mathematica* notebook [44], as shown in Figure 17**A**. This image compared to the given histological slide (Figure 17**B**), and any cluster identified as corresponding to an MPPZ in this comparison was highlighted in the reconstruction.

### Generating the reconstruction

The final step in the processing pipeline is to generate a volume with evenly-spaced voxels. Thus far the volumes have been given by the sequence of slides, which have an uneven separation as a result of some slides being discarded. This issue was solved by using linear interpolation to fill the space between slides with separation greater than 10 *μ*m. Let {I*_n_*}*_n_*_∊*N*_ be the sequence of images representing slides, where *N* is an increasing sequence of integers representing the position of the slides in the tissue: if two slides have positions *n*_1_ and *n*_2_, then the physical distance between them in the tissue sample is 5|*n*_1_ − *n*_2_| *μ*m. The aim is to generate a sequence {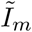} of images representing 10 *μ*m slices through the volume. Let 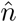 be the maximum value in *N*, and for an integer 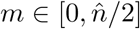, define

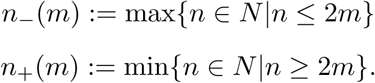

This allows us to define 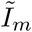:

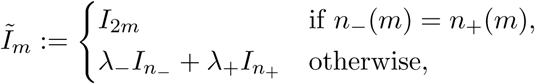

where

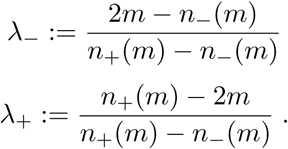

This stack of images represents a volume with uniform voxel length of 10 *μ*m, where each image is either an image from the original stack or a weighted average of two slides in the stack. The resulting volume provides a homogenised visual representation of the three-dimensional microarchitecture of the myometrium, which can be used to identify MPPZs and bridges between longitudinal and circular myometrium.

To determine if bridges were co-located with vasculature, each bridge within the histological slide was scored accordingly. Any bridges absent of adjacent vasculature within the given histological slide were tracked through the slides, guided by the 3-dimensional reconstruction, to exhaustively inspect for the presence of vasculature. The proposed paths of excitation marked out in Fig. 5 were identified as follows. For each MPPZ in close proximity to an identified initiation event, the histological slides were examined to identify the point at which the MPPZ attached to the longitudinal myometrium. The bundle at this location was tagged in the reconstructed tissue and bundles were tagged in slides working outward from this initial slide in a recursive manner: for the next slide in the stack, the tagged points in the previous slide were marked in the reconstruction. If these marked points corresponded to myometrial smooth muscle in the histological slide, the bundles containing marked points were noted. Otherwise, the recursion terminated. High-resolution representations of these reconstructions were used to visualise the connectivity in the myometrium and the pacemaker zones, as detailed in the supporting information. All three-dimensional reconstructions were visualised using 3D Slicer [12] (www.slicer.org).

## Acknowledgments

EJL, AMB, and HAB were supported by a grant from the Medical Research Council (United Kingdom), grant number G0900208-2. The funders had no role in study design, data collection and analysis, decision to publish, or preparation of the manuscript.

## References

1. J. Atia, C. McCloskey, A. S. Shmygol, D. A. Rand, H. A. van den Berg, and A. M. Blanks. Reconstruction of cell surface densities of ion pumps, exchangers, and channels from mRNA expression, conductance kinetics, whole-cell calcium, and current-clamp voltage recordings, with an application to human uterine smooth muscle cells. PLoS Comput Biol, 12(4):e1004828, 2016.

2. J. Bonventre, Z. Huang, M. Taheri, E. O’Leary, E. Li, and A. Moskowitz, M.A. Sapirstein. Reduced fertility and postischaemic brain injury in mice deficient in cytosolic phospholipase A2. Nature, 390(6660):622–625, 1997.

3. A. Brainard, V. Korovkina, and S. England. Potassium channels and uterine function. Semin Cell Dev Biol, 18(3):332–339, 2007.

4. J. R. Brody and G. R. Cunha. Histologic, morphometric, and immunocytochemical analysis of myometrial development in rats and mice: I. Normal development. Am J Anat, 186:1–20, 1989.

5. R. L. Burden and J. D. Faires. Numerical Analysis. Brooks/Cole, Cengage Learning, Belmont, CA, USA, 2005.

6. R. Casteels and H. Kuriyama. Membrane potential and ionic content in pregnant and non-pregnant rat myometrium. J Physiol, 177:263–287, 1965.

7. J. Cha, A. Bartos, M. Egashira, H. Haraguchi, T. Saito-Fujita, E. Leishman, H. Bradshaw, S. K. Dey, and Y. Hirota. Combinatory approaches prevent preterm birth profoundly exacerbated by gene-environment interactions. J Clin Invest, 123(9):4063–4075, 2013.

8. J. Cha, A. Bartos, C. Park, X. Sun, Y. Li, S. Cha, R. Ajima, T. Ho, H.Y. Yamaguchi, and S. Dey. Appropriate crypt formation in the uterus for embryo homing and implantation requires Wnt5a-ROR signaling. Cell Rep, 8(2):382–392, 2014.

9. J. C. Condon, P. Jeyasuria, J. M. Faust, and C. R. Mendelson. Surfactant protein secreted by the maturing mouse fetal lung acts as a hormone that signals the initiation of parturition. Proc Natl Acad Sci U S A, 101(14):4978–4983, 2004.

10. J. C. Condon, P. Jeyasuria, J. M. Faust, J. W. Wilson, and C. R. Mendelson. A decline in the levels of progesterone receptor coactivators in the pregnant uterus at term may antagonize progesterone receptor function and contribute to the initiation of parturition. Proc Natl Acad Sci U S A, 100(16):9518–9523, 2003.

11. T. Daikoku, J. Cha, X. Sun, S. Tranguch, H. Xie, T. Fujita, Y. Hirota, J. Lydon, F. DeMayo, R. Maxson, and S. Dey. Conditional deletion of Msx homeobox genes in the uterus inhibits blastocyst implantation by altering uterine receptivity. Dev Cell, 21(6):1014–1025, 2011.

12. A. Fedorov, R. Beichel, J. Kalpathy-Cramer, J. Finet, J. C. Fillion-Robin, S. Pujol, C. Bauer, D. Jennings, F. Fennessy, M. Sonka, J. Buatti, S. Aylward, J. V. Miller, S. Pieper, and R. Kikinis. 3D Slicer as an image computing platform for the quantitative imaging network. Magn Reson Imaging, 30(9):1323–41, 2012.

13. A. Fuchs, F. Fuchs, P. Husslein, M. Soloff, and M. Fernström. Oxytocin receptors and human parturition: a dual role for oxytocin in the initiation of labor. Science, 215(4538):1396–1398, 1982.

14. A. Fuchs, S. Periyasamy, M. Alexandrova, and M. Soloff. Correlation between oxytocin receptor concentration and responsiveness to oxytocin in pregnant rat myometrium: effects of ovarian steroids. Endocrinology, 113(2):742–749, 1983.

15. R. E. Garfield, M. G. Blennerhassett, and S. M. Miller. Control of myometrial contractility: role and regulation of gap junctions. Oxf Rev Reprod Biol, 10:436–490, 1988.

16. R. E. Garfield and W. L. Maner. Physiology and electrical activity of uterine contractions. Semin Cell Dev Biol, 18:289–295, 2007.

17. R. E. Garfield, S. Sims, and E. E. Daniel. Gap junctions: their presence and necessity in myometrium during parturition. Science, 198(4320):958–960, 1977.

18. J. M. Geusebroek, A. W. M. Smeulders, and J. Van De Weijer. Fast anisotropic gauss filtering. IEEE Transactions on Image Processing, 12(8):938–943, 2003.

19. J. J. Goldberger and J. Ng. Practical Signal and Image Processing in Clinical Cardiology. Springer Science & Business Media, 1^st^ edition, 2010.

20. I. Greenwood, S. Yeung, R. Tribe, and S. Ohya. Loss of functional K^+^ channels encoded by ether-’a-go-go-related genes in mouse myometrium prior to labour onset. J Physiol, 587(10):2313–2326, 2009.

21. G. Gross, T. Imamura, C. Luedke, S. Vogt, L. Olson, Y. Nelson, D.M. Sadovsky, and L. Muglia. Opposing actions of prostaglandins and oxytocin determine the onset of murine labor. Proc Natl Acad Sci U S A, 95(20):11875–11879, 1998.

22. F. T. Hammad, B. Stephen, L. Lubbad, J. F. Morrison, and W. J. Lammers. Macroscopic electrical propagation in the guinea pig urinary bladder. Am J Physiol Renal Physiol, 307(F172–F182), 2014.

23. Y. Hirota, J. Cha, M. Yoshie, T. Daikoku, and S. K. Dey. Heightened uterine mammalian target of rapamycin complex 1 (mTORC1) signaling provokes preterm birth in mice. Proc Natl Acad Sci U S A, 108(44):18073–8, 2011.

24. J. Keener and J. Sneyd. Mathematical Physiology. Springer-Verlag, New York, USA, 2004.

25. R. Khan, S. Smith, J. Morrison, and M. Ashford. Properties of large-conductance K^+^ channels in human myometrium during pregnancy and labour. Proc Biol Sci, 251(1330):9–15, 1993.

26. G. Knock, S. Smirnov, and P. Aaronson. Voltage-gated K^+^ currents in freshly isolated myocytes of the pregnant human myometrium. J Physiol, 518(3):769–781, 1999.

27. W. Lammers and B. Stephen. Origin and propagation of individual slow waves along the intact feline small intestine. Exp Physiol, 93:334–346, 2008.

28. W. J. Lammers, B. Stephen, M. A. Al-Sultan, S. B. Subramanya, and A. M. Blanks. The location of pacemakers in the uteri of pregnant guinea pigs and rats. Am J Physiol Regul Integr Comp Physiol, 309(11):R1439–R1446, 2015.

29. W. J. Lammers, L. Ver Donck, B. Stephen, D. Smets, and J. A. Schuurkes. Origin and propagation of the slow wave in the canine stomach: outline of the gastric conduction system. Am J Physiol Gastrointest Liver Physiol, 296(G1200–G1210), 2009.

30. E. J. Lutton, W. J. E. P. Lammers, S. James, H. A. van den Berg, and A. M. Blanks. A computational method for three-dimensional reconstruction of the microarchitecture of myometrial smooth muscle from histological sections. PLoS ONE, 12(3):e0173404, 2017.

31. J. Marshall. Regulation of activity in uterine smooth muscle. Physiological Reviews Supplement, 5:213–227, 1962.

32. C. McCloskey, C. Rada, E. Bailey, S. McCavera, H. van den Berg, J. Atia, D. Rand, A. Shmygol, Y. Chan, J. Quenby, S. Brosens, M. Vatish, J. Zhang, J. Denton, M. Taggart, C. Kettleborough, D. Tickle, J. Jerman, P. Wright, T. Dale, S. Kanumilli, D. Trezise, S. Thornton, P. Brown, R. Catalano, N. Lin, S. England, and A. Blanks. The inwardly rectifying K^+^ channel KIR7.1 controls uterine excitability throughout pregnancy. EMBO Mol Med, 6(9):1161–1174, 2014.

33. N. Otsu. A threshold selection method from gray-level histograms. IEEE transactions on systems, man, and cybernetics, 9(1):62–66, 1979.

34. H. Parkington, J. Stevenson, M. Tonta, J. Paul, T. Butler, K. Maiti, E. Chan, P. Sheehan, S. Brennecke, H. Coleman, and R. Smith. Diminished hERG K^+^ channel activity facilitates strong human labour contractions but is dysregulated in obese women. Nat Commun, 5(4108), 2014.

35. H. Parkington, M. Tonta, S. Brennecke, and H. Coleman. Contractile activity, membrane potential, and cytoplasmic calcium in human uterine smooth muscle in the third trimester of pregnancy and during labor. Am J Obstet Gynecol, 181(6):1445–1451, 1999.

36. H. Parkington, M. Tonta, N. Davies, S. Brennecke, and H. Coleman. Hyperpolarization and slowing of the rate of contraction in human uterus in pregnancy by prostaglandins E_2_ and F_2α:_ involvement of the Na^+^ pump. J Physiol, 514(Pt 1):229–243, 1999.

37. S. Pierce, J. Kresowik, K. Lamping, and S. England. Overexpression of SK3 channels dampens uterine contractility to prevent preterm labor in mice. Biol Reprod, 78(6):1058–1063, 2008.

38. C. Rabotti and M. Mischi. Propagation of electrical activity in uterine muscle during pregnancy: A review. Acta Physiologica, 213(2):406–416, 2014.

39. N. E. Renthal, C. C. Chen, K. C. Williams, R. D. Gerard, J. Prange-Kiel, and C. R. Mendelson. miR-200 family and targets, ZEB1 and ZEB2, modulate uterine quiescence and contractility during pregnancy and labor. Proc Natl Acad Sci U S A, 107(48):20828–33, 2010.

40. H. Song, H. Lim, B. Paria, H. Matsumoto, L. Swift, J. Morrow, J. Bonventre, and S. Dey. Cytosolic phospholipase A2alpha is crucial [correction of A2alpha deficiency is crucial] for ‘on-time’ embryo implantation that directs subsequent development. Development, 129(12):2879–2889, 2002.

41. Y. Sugimoto, A. Yamasaki, E. Segi, K. Tsuboi, Y. Aze, T. Nishimura, H. Oida, N. Yoshida, T. Tanaka, M. Katsuyama, K. Hasumoto, T. Murata, M. Hirata, F. Ushikubi, M. Negishi, A. Ichikawa, and S. Narumiya. Failure of parturition in mice lacking the prostaglandin F receptor. Science, 277(5326):681–683, 1997.

42. X. Sun, L. Zhang, H. Xie, H. Wan, B. Magella, J. Whitsett, and S. Dey. Kruppel-like factor 5 (KLF5) is critical for conferring uterine receptivity to implantation. Proc Natl Acad Sci U S A, 109(4):1145–1150, 2012.

43. K. D. Toennies. Guide to Medical Image Analysis. Springer Berlin Heidelberg, Berlin, Germany, 2012.

44. Wolfram Research, Inc. Mathematica 8.0.

45. X. Ye, K. Hama, J. Contos, B. Anliker, A. Inoue, M. Skinner, H. Suzuki, T. Amano, G. Kennedy, H. Arai, J. Aoki, and J. Chun. LPA3-mediated lysophosphatidic acid signalling in embryo implantation and spacing. Nature, 435(7038):104–108, 2005.

46. J. Yuan, J. Cha, W. Deng, A. Bartos, X. Sun, H. Ho, J. Borg, T. Yamaguchi, Y. Yang, and S. Dey. Planar cell polarity signaling in the uterus directs appropriate positioning of the crypt for embryo implantation. Proc Natl Acad Sci U S A, 113(50):E8079–E8088, 2016.

